# Temporal Expression Prediction by Integrating Genome Dynamics via Spatio-temporal GNNs

**DOI:** 10.1101/2025.10.07.678022

**Authors:** Beyza Kaya, Emre Sefer

## Abstract

Temporal gene expression is being analyzed via high-throughput profiling of molecular data over time. The expression values of genes are impacted by their previous expression values as well as the expression of interacting genes over time. Hi-C provides us with a broad genome-wide perspective on the interacting dynamics of genes. In this paper, we propose neural network-based spatio-temporal graph approaches STEPmr and STEPmi to predict changes in mRNA and miRNA expression over time, respectively. Both approaches can integrate a diverse set of Hi-C datasets and features obtained from Hi-C when predicting temporal expression patterns.

Our methods can predict mRNA and miRNA expression with 77% and 93% correlation and ith mean squared errors of 0.21 and 0.01, explaining 59.1% and 88% of the variance, respectively. Important characteristics of the genes with the highest performances in both datasets are that they are structural signaling genes or transcriptional regulators involved in fundamental processes such as homeostasis, development, and RNA processing. Additionally, they are not limited to a specific cell type, but rather show constant expression throughout different tissues. In contrast, the lowest-performed genes generally behave in context-dependent expression patterns, where they include condition-specific biological functions instead of vital biological activities. These findings suggest a model of gene regulation and its predictability that is impacted by interacting gene dynamics. Our code and datasets are publicly available at https://github.com/seferlab/temporalhic.

## 1. Introduction

Time series studies are widely employed to investigate numerous biological phenomena [1, 2]. Understanding the intricate biological processes of development [3], differentiation [4], and disease progression via immune system activity and cellular stress responses [5, 6] requires a knowledge of the dynamic regulation of gene expression throughout time. In fact, an analysis of the Gene Expression Omnibus (GEO), the most extensive public repository of gene expression data, revealed that approximately one-third of the archived datasets are derived from experiments designed to capture biological changes over time [7]. Both messenger RNAs (mRNAs) and microRNAs (miRNAs) are crucial for these actions; mRNAs provide blueprints for protein synthesis [8], while miRNAs function as post-transcriptional regulators of gene expression [9]. Predicting gene expression levels accurately over a range of time points can reveal important information about the underlying biochemical processes and regulatory relationships [10, 11].

There are a number of challenges with obtaining miRNA Expression datasets, so collecting a high-resolution dataset over dense time intervals is costly. Some of these challenges are smaller miRNA size [12], low abundance, sequence similarity, and the maturation process [13]. miRNAs are very small (typically 18-24 nucleotides) compared to mRNAs (hundreds to thousands of nucleotides) [14]. Some miRNAs are expressed at very low levels, making their detection and quantification more challenging than often more abundant mRNAs. miRNAs often have highly similar sequences within families, making it difficult to distinguish individual family members without highly specific assays. miRNAs undergo a complex maturation process, from primary to precursor to mature miRNA. Measuring the mature, active form can be more complex than simply measuring a full-length mRNA transcript. Similarly, collecting a high-resolution mRNA dataset over dense time intervals is also costly. Some of the challenges associated with mRNA experiments are a larger transcriptome, splicing variants, and degradation. The mRNA transcriptome is much larger and more diverse than the miRNA transcriptome, meaning comprehensive analysis often requires higher sequencing depth or broader microarray coverage. Many genes produce multiple mRNA splicing variants [15], which can complicate analysis if you need to distinguish between them. mRNA is generally less stable than miRNA, requiring careful handling to prevent degradation during sample preparation [16].

Although mRNA expression data have traditionally served as the main source for high-throughput time series analysis, a growing number of other genomic regulatory elements are now being examined across time. These include time-resolved measurements of miRNA levels [17], ChIP-Seq experiments used to map transcription factor binding sites [18], as well as various epigenetic markers such as DNA methylation [19] and histone modifications [20]. With advancements in high-throughput technologies, researchers are increasingly conducting experiments that track multiple such regulatory features simultaneously [18, 21], enabling the integration of diverse datasets to gain deeper insights into cellular processes.

Moreover, Hi-C datasets play a crucial role in improving gene expression prediction by capturing the three-dimensional (3D) architecture of the genome [22, 23]. Unlike traditional linear genomic data, Hi-C provides genome-wide maps of chromatin interactions, revealing how distal regulatory elements such as enhancers physically contact gene promoters within the nuclear space [24]. This spatial organization is essential for accurate gene regulation, as gene expression is not solely determined by sequence proximity but also by 3D genomic context. Integrating Hi-C data with epigenomic and transcriptomic information enables more precise modeling of gene regulatory networks and enhances our ability to predict expression patterns across different cell types and conditions [25, 26, 27]. Consequently, Hi-C datasets offer critical insight into long-range regulatory mechanisms [28] and help uncover non-coding functional regions that influence transcriptional activity [29].

In this paper, we focus on two aims: 1-Enhancing the quality of temporal gene expression prediction for both miRNAs and mRNAs by integrating the topological and spatial organization of the genome as revealed by Hi-C contact maps, and 2-Analyzing the output of temporal expression predictions and performances in a biological context such as in terms of pathway analysis, genome shape analysis, functional analysis. Our first objective is to utilize biological structural features obtained from Hi-C data, as well as gene-gene interaction data, to estimate the expression dynamics of target genes across time by using deep learning. We address this problem through the use of graph-based modeling via Graph Neural Networks (GNNs) [30], in which the edges are defined by spatial or regulatory interactions, and the genes are represented as nodes. We construct dynamic graph representations for every time point, with the nodes representing target genes of either mRNAs or miRNAs, in order to better represent the genomic context. Hi-C contact frequencies, which are mapped to their appropriate chromosomal locations to match genuine expression data across the time series, are used to create edges between nodes. TAD boundaries, A/B chromatin compartments, insulation scores, and the actual expression levels of the associated genes are among the biologically relevant features that are used to weigh each edge. Via such a weighted combination of a distinct set of biologically relevant features, the resulting graph is composed of a richer and more biologically significant structure by encoding both the functional relevance of gene interactions and spatial proximity from chromatin conformation.

Based on spatio-temporal graph convolutional networks, we developed two distinct neural architectures, STEPmr (**S**patio-**T**emporal **E**xpression **P**rediction for **mR**NA) and STEPmi (**S**patio-**T**emporal **E**xpression **P**rediction for **mi**RNA) for mRNA and miRNA expression prediction, respectively, in order to reflect the intricate spatio-temporal dynamics of gene regulation. STEPmr is better at handling sparser temporal datasets and a limited number of nodes (target genes) in Graph Neural Networks, so we use it for mRNA expression prediction. While using spatio-temporal graph neural networks, we use Node2vec-based [31] embeddings to enhance node representations in addition to edge-level features. We generate temporal Node2vec embeddings for every gene at every time point, in contrast to conventional static embeddings, which enables the model to reflect dynamic changes in each gene’s neighborhood structure over time.

STEPmr consists of two spatio-temporal (ST) blocks followed by an attention-based aggregation module. The aggregated output is passed through a temporal convolution layer, a set of linear transformation layers, and an expression projection block that includes layer normalization, generating a single-step prediction using a sliding window approach. On the other hand, STEPmi uses three ST blocks, a four-layer bidirectional LSTM, and a four-head multi-head attention mechanism to handle the wider and more complex structure of miRNA expression data. The same output and expression projection blocks that are used in STEPmr are then applied to the miRNA predictions. Both STEPmi and STEPmr are made to capture the distinct temporal and structural characteristics of their respective datasets since miRNA and mRNA gene expressions show distinct temporal patterns. Whereas STEPmi is designed for the more complex miRNA data, which has a greater number of target genes and time points, STEPmr efficiently manages the sparse nature of mRNA data.

For our second aim, we use Gene Ontology enrichment across genes categorized by their expression slopes (upregulated, stable, or downregulated) to examine presented temporal expression patterns in a biological context. Along with possible indicators of inflammatory conditions, this research identified important biological processes for target genes of miRNA and mRNA such as neural remodeling, changed cell communication, metabolic reprogramming, and increased synthetic ability. These patterns show that specialized neural functions are giving way to more generic cellular operations like energy consumption and protein synthesis, which are signs of neuroplastic adaptation or cellular reaction to environmental stress. The significance and possible implications of precise temporal gene expression modeling in developmental, stress-related, or neurodegenerative disorders are highlighted by such biological interpretations.

Overall, in addition to being the first approaches to predict temporal gene expression values via spatio-temporal GNNs [32, 33], our approaches introduce a number of novel methodological components that extend the existing approaches, which can be summarized as:

- **Spatio-temporal GNNs:** We use GNNs, especially spatio-temporal GNNs, for the first time to predict temporal gene expression values, which can model the interactions between multiple genes.
- **Temporal Node2vec Embeddings:** Alternative to using static node embeddings, we generate temporal Node2Vec representations for every gene.These embeddings capture the dynamic neighborhood structure of each gene across time by calculating the cosine similarity between time points.
- **Biologically-Informed Edge Weights:** We use Hi-C interactions to model the dependencies and the impacts between a distinct set of target genes over time. Also, Hi-C contact frequencies are not the only basis for graph edges. We enhance them by incorporating: 1-Real-time gene expression values of interacting genes at each time point, 2-Structural genomic features such as TAD boundaries, A/B compartments, and insulation scores. This produces edge weights that are dynamic and biologically significant, reflecting the transcriptional process as well as the spatial organization of the genome.
- **Customized Multi-Component Loss Function:** Instead of using the conventional mean squared error (MSE) to handle direct convergence, we present a composite loss function made up of: 1-Directional loss which penalizes incorrect expression direction (up/down), 2-Temporal trend loss which enforces consistency with local temporal patterns, 3-Prediction consistency loss, which reduces noisy fluctuations in prediction. This composite loss facilitates robust training and captures fine-grained expression dynamics.

When combined, these developments provide an extensive and adaptable method for simulating the dynamics of gene expression across spatial organization, time, and gene-specific interactions.

### 1.1. Related Work

Gene expression prediction, estimating the level of gene activity (expression) under specific biological conditions, has been mostly studied from a static point of view. Early methods employed statistical approaches using microarray or RNA-seq data to correlate expression levels with cis-regulatory elements [34]. However, recent advancements have shifted toward machine learning and deep learning models, which can capture nonlinear relationships and integrate heterogeneous data sources to improve prediction accuracy [35, 36]. Machine learning and deep learning algorithms have become increasingly prominent in gene expression analysis due to their inherent capacity to discern complex patterns within large, high-dimensional datasets. This ability to identify intricate relationships is a fundamental advantage over more rudimentary statistical methods, allowing for a move beyond simple correlations to a more predictive and mechanistic understanding, particularly given the combinatorial nature of gene regulation. Deep learning models, such as convolutional neural networks (CNNs) and transformer-based architectures, have shown particular promise in modeling gene expression from raw genomic sequences. For instance, models like DeepSEA [37] and Basenji [38] use DNA sequence data to predict expression and chromatin features at high resolution, while more recent transformer-based models like Enformer have further improved long-range regulatory element modeling [27]. Another set of methods [39, 40, 41, 42] predict gene expression and interactions by using histone modifications and other epigenetic marks. Another prominent example of a deep learning application is D-GEX [43], a multi-task multilayer feedforward neural network designed to infer the expression of target genes from a smaller, carefully selected set of approximately 1000 landmark genes. D-GEX has demonstrated significant performance improvements over traditional linear regression. The emergence of “hub units” in D-GEX, which capture strong local correlations, illustrates how these models learn abstract, biologically relevant features. These tools enable researchers to investigate how genetic variants influence expression and contribute to complex traits or diseases. Despite these advances, challenges remain, including data sparsity across tissues, interpretability of models, and generalizability across species. Ongoing efforts focus on integrating multi-omics data and developing interpretable models to enhance biological insight and translational applications.

Among the techniques that consider interactions in expression prediction, none of them has focused on dynamic spatio-temporal prediction. For instance, [26] integrates Hi-C data with epigenomic and transcriptomic information, enabling more precise modeling of gene regulatory networks and enhancing the static expression prediction performance. [27] predicts static gene expression by integrating sequence and long-range interactions. Similarly, [25] focuses on inferring static gene co-expression from chromatin contacts with a graph attention network. Lastly, [44] proposes DeepCBA, which can take the effect of chromatin interactions into account on static target gene expression prediction.

Another set of studies considers the gene expression prediction problem from a more dynamic point of view, as a spatio-temporal prediction problem. [45] introduces a gene expression prediction model called L-GEPM, which is built on long short-term memory (LSTM) neural networks. This model is designed to capture complex, nonlinear patterns that influence gene expression and leverages these learned patterns to forecast the expression levels of specific target genes. [1, 2] utilized statistical time-series decomposition techniques to predict temporal gene expression. However, neither of these approaches can take into account the interactions or dependencies between multiple genes. Similarly, [46] predicts time-series transcriptomic gene expression by combining LSTM with Empirical Mode Decomposition. [47] predicts time series gene expression via Recurrent Neural Networks, without modeling the interactions between genes. Even though GNNs have recently become quite popular [48, 30], they have not been utilized in temporal gene expression prediction. Among GNNS, spatio-temporal GNNs [32, 49] have previously been applied to a distinct set of domains ranging from PM_2.5_ concentration prediction [33] to traffic prediction [50, 51], with successful results. However, this is the first study to adapt them to temporal gene expression prediction.

## 2. Materials and Methods

### 2.1. Dataset Preparation

We use distinct datasets for mRNA and miRNA expression due to the variations in their temporal profiles, target genes, and interaction patterns. These datasets are also used in [1], without being made publicly available. Here, we make these biological experiment datasets publicly available as well.

Both datasets contain expression values across multiple time steps. While the miRNA dataset has 162 genes and 159 time points, the mRNA dataset only has 49 genes and 43 time points after mapping miRNA families to their associated target genes. We integrate these expression datasets with Hi-C contact matrices by mapping gene names to chromosome numbers and positions. Additionally, we incorporate supplementary biological features into the mapped Hi-C and mRNA/miRNA dataset to enhance the representation.

#### 2.1.1 miRNA Expression Data and Hi-C Integration

We used a time-series expression dataset with miRNA family-level expression values labeled with MIMAT accession IDs. We used a multi-stage mapping strategy, utilizing multiple publicly accessible databases, to link each miRNA family to its associated target genes. This ensured extensive coverage and prevented the elimination of unmapped families. First, we used both conserved and non-conserved miRNA family annotations specific to Mus Musculus, matched by MIMAT IDs, to query the TargetScanMouse database [52]. We chose the target gene with the highest prediction score from the miRDB database [53] for miRNA families that were not present in the TargetScanMouse database. Lastly, we searched the ViRBase v3.0 database [54] using miRBase accession numbers for any unmapped families that remained, limiting our search to Mus musculus and all detection methods.

We integrated target gene information with Hi-C interaction data at a 1 Mb resolution. We matched the mapped genes with chromosome-wise Hi-C contact maps from mESC datasets in order to incorporate spatial genome organization. The start coordinate of each gene was translated to the closest Hi-C bin center:

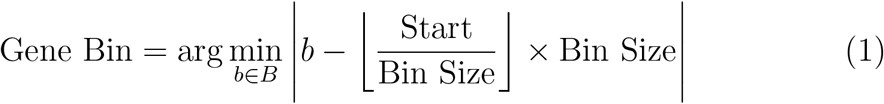

where *B* is the set of available Hi-C bins on the corresponding chromosome, partitioned into 1 Mb regions. This enabled genes whose start locations did not precisely coincide with bin borders to be matched to Hi-C bins based on proximity. Following, the associated Hi-C interaction values were extracted by building paired gene combinations.

We added A/B compartments, insulation scores, and topologically associating domain (TAD) boundaries to the gene interaction dataset in order to enhance it with chromatin-level context. These characteristics were obtained by processing normalized Hi-C interaction matrices for each chromosome separately at a resolution of 1 Mb. In accordance with Lieberman-Aiden et al. [55], we used the sign of the first eigenvector of the Pearson correlation matrix derived from the Hi-C data to allocate genomic bins to either the A (active) or B (inactive) compartments. In particular, we calculated the correlation matrix *C* and carried out eigenvector decomposition, beginning with the symmetric interaction matrix *H*^(*c*)^ ∈ ℝ^n×n^ for chromosome *c*, where *H*_*ij*_ indicates the contact frequency between bins *i* and *j*. Bins were assigned to compartment A if they had a positive entry in the leading eigenvector *v* ∈ ℝ^n^; if not, they were assigned to compartment B.

We utilized an insulating score-based approach, adapted from Crane et al. [56], to determine TAD boundaries. The insulation score *S*_*i*_ was obtained by calculating the average Hi-C interaction for each bin *i* inside a fixed-size square submatrix centered on the diagonal. We used the z-score to normalize these scores:

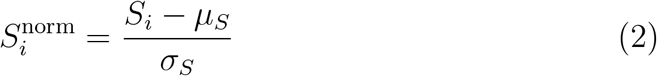

where *µ*_*S*_ and *σ*_*S*_ are the mean and standard deviation of scores, respectively. Possible TAD boundaries were then determined by applying peak-finding methods on the negative of the aforementioned insulation profile in order to detect local minima in 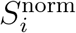 . We annotated the following for each mapped gene-gene pair: 1The locus’s chromatin compartment; 2-The normalized insulation score; and 3-The shortest distance (in bins) to the closest TAD boundary.

Every gene in the expression dataset interacts with other genes at different times, producing varying expression values for the same gene at the same time. We calculated the mean expression values for each gene at each time point in order to produce stable and consistent representations of gene expression for the same gene across various interactions at each time point. Each gene’s expression values at a particular time point precisely represented its overall interaction profile in this way. Since the same node (gene) would be linked to several expression values without the use of this accumulation, the resulting graph would not be consistent across time steps, making it more difficult to create a completely connected and unified graph representation. Let 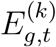 represent gene *g*’s expression value at time *t* as determined by its *k*-th interaction. We calculate the mean expression value 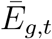 as follows if gene *g* has *N*_*g,t*_ interactions at time t:

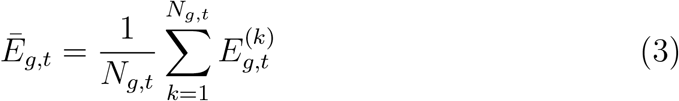

#### 2.1.2. mRNA Expression Data and Hi-C Integration

We used gene expression data obtained for 49 target genes at 43 different time points for the mRNA dataset. To confirm that gene identities are accurate, we retrieved the most recent gene names via the Ensembl REST API. The exact genomic start and end locations of every gene were determined, and they were mapped to the corresponding chromosomes. We used the same preprocessing technique that was employed to create the miRNA dataset after establishing reliable gene names. This involved merging Hi-C chromatin interaction data and mapping genes to genomic bins and additional biological features. Achieving an extensive representation for the mRNA dataset is more important than the miRNA dataset because of the mRNA dataset’s intrinsically small number of genes (49), as well as its limited time periods. To compensate for the lack of raw temporal coverage in such a small dataset, it was necessary to include further biological background, such as Hi-C interactions, A/B compartments, insulation scores, and TAD boundaries. Each sample consists of a pair of interacting genes that are distinguished by their Hi-C interaction frequency, time-specific expression profiles, and other genomic characteristics like compartment status, insulation scores, and distances to TAD boundaries, as in the miRNA dataset.

### 2.2. Proposed Architectures: STEPmr and STEPmi

As summarized in Figure 1, we propose a spatio-temporal graph-based framework that aims at predicting mRNA/miRNA expression values (STEPmr, STEPmi) across multiple time points while preserving the biology-based network of gene interactions constructed from genomic features. The objective is to ensure accurate prediction of dynamic gene expressions, with targeted optimization of Pearson and Spearman correlation measures to maintain relationships of gene interactions.

**Figure 1.**
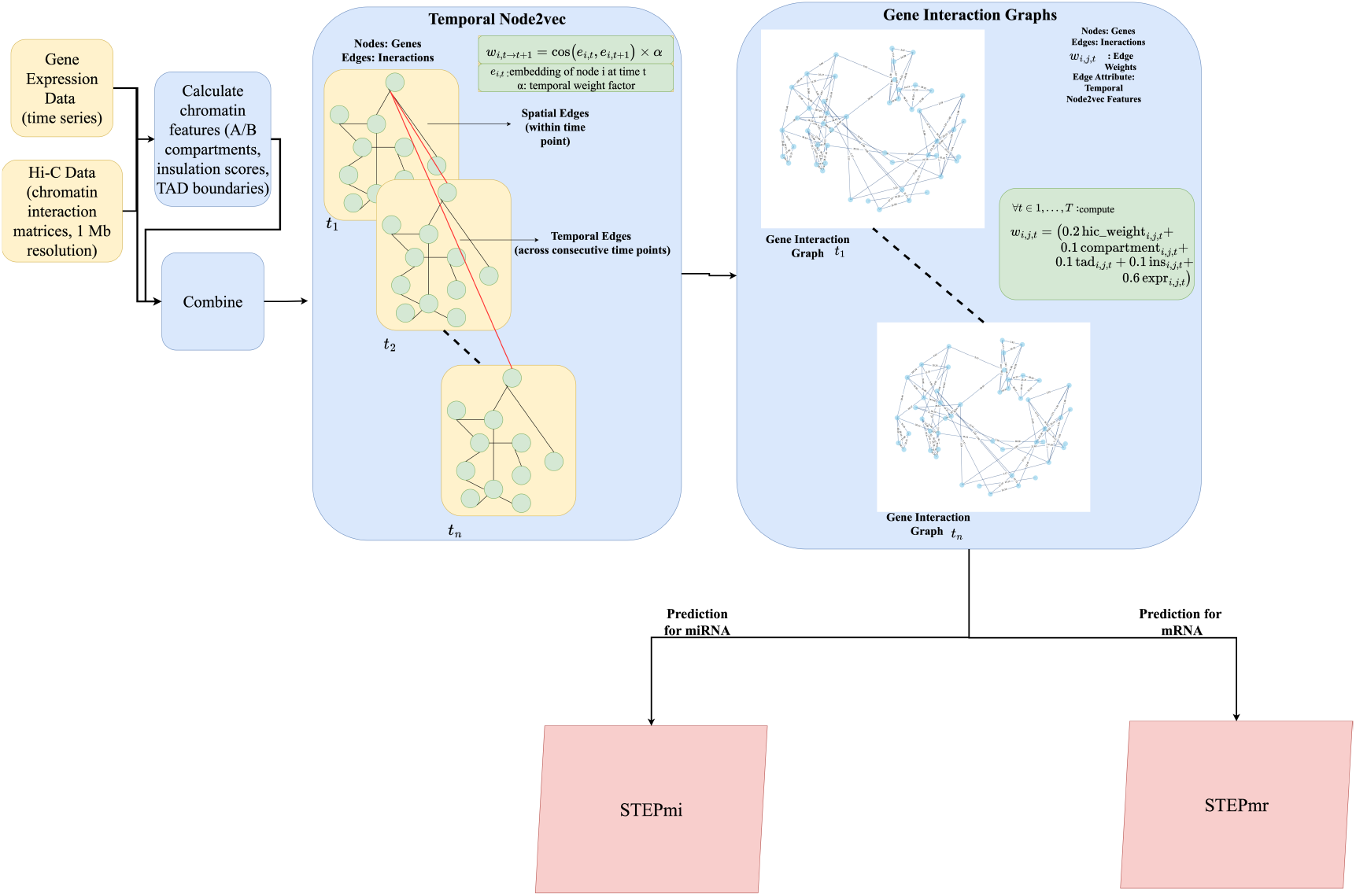
Overview of the proposed pipeline for predicting temporal gene expression from multiple datasets, including historical expression values.

We first created an interaction graph where each node corresponds to a gene, while the edges are a weighted sum of biological features (see Section 2.1). Using the constructed interaction graph, we implement a temporal Node2vec approach to capture the temporal dynamics of gene interactions. Then, using the interaction graph and temporal Node2vec embeddings, we created input sequences of varying window sizes from the time-series expression data to capture temporal dependencies. The generated sequences were used to train a spatio-temporal graph convolutional network (STGCN), enhanced with different components such as gene-based attention, Multi-Head Attention, and bidirectional LSTM.

We integrated context-aware representation modules into our graph blocks to tackle issues with over-smoothing, a scenario where node representations become indistinguishable across layers, especially in dense graphs or when edge weights are too close. This problem is particularly noticeable when graph formation involves delicate or intricate mappings from raw data, as discussed in [57]. Such behavior can reduce predictive power by causing model outputs to collapse toward the mean. Our goal is to improve learning stability across layers while maintaining different node representations.

### 2.3. Temporal Graph Construction

Genes were represented as nodes in the gene interaction graph, and edge weights were defined using a variety of biological characteristics obtained from genomic and epigenomic data, with specific features such as:

- **Hi-C interaction frequencies**, which measure how often genomic loci come into contact, revealing their three-dimensional spatial relationships.

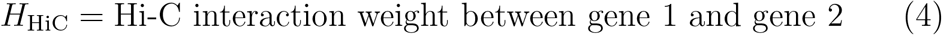
- **TAD boundary distances**, capturing how close genes are to TAD boundaries, with genes in proximity often influenced by similar gene expression dynamics. The *H*_TAD_ weight is calculated as:

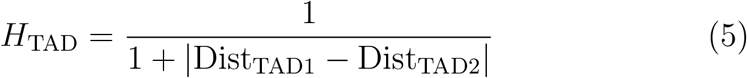

where Dist_TAD1_ and Dist_TAD2_ are the distances of genes 1 and 2 from their respective TAD boundaries.
- **Insulation scores**, indicating the degree of chromatin insulation between regions. The *H*_Ins_ weight for edge weight calculation is:

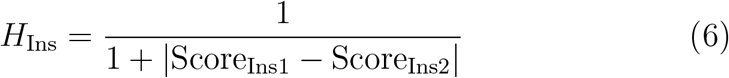

where Score_Ins1_ and Score_Ins2_ are the insulation scores of genes 1 and 2.
- **Compartment similarity**, serving as an indicator of whether genes reside in the same chromatin compartment (A/B). The *H*_Comp_ is calculated as:

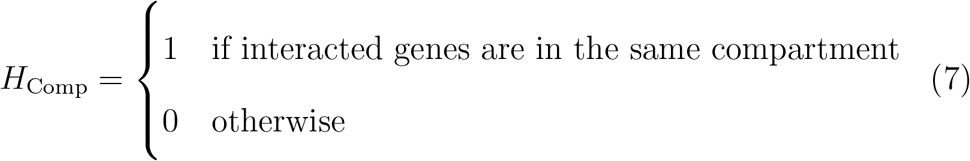
- **Gene Expression similarity**, captures how similar the expression levels of interacted genes at a given time point. The *H*_Expr_ is calculated as:

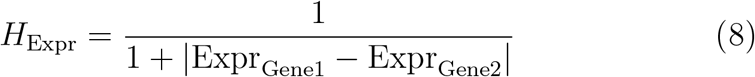

where Expr_Gene1_ and Expr_Gene2_ are the normalized expression values of genes 1 and 2 at a given time point.

Then the edge weight become:

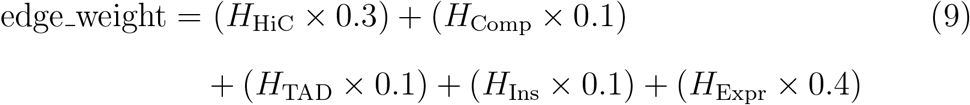

We constructed an undirected graph for each gene pair in the dataset using the calculated edge weights. For each time point, a new graph was constructed with updated edge weights based on the normalized expression values of interacting genes at that time.

### 2.4. Temporal Node2vec Embeddings

The goal of Temporal Node2vec is to establish node embeddings that encapsulate the dynamic behavior of genes across time as well as their static spatial relationships at each time point. To achieve this, the traditional Node2vec algorithm [31], which typically conducts random walks on a single, static graph, is modified to work on a temporal graph in which nodes are capable of evolving over time. The core components of the temporal Node2vec model are:

- **Spatial Embeddings:** These are embeddings learned for nodes at each individual time point.
- **Temporal Edges:** Temporal edges connect nodes across adjacent time points, where the edge weight is based on the similarity of node embeddings between time points, allowing the model to capture the progression of gene interactions across time. The weight of the temporal edge connecting two nodes *u*_*t*_ and *u*_*t′*_ (representing the same gene at two different time points *t* and *t*^*′*^) is calculated as:

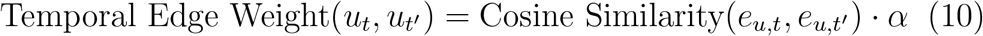

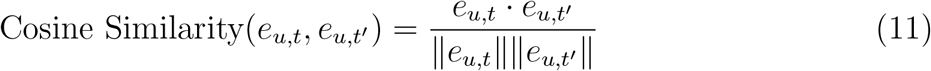

where *u*_*t*_ and *u*_*t′*_ represent the nodes for gene *u* at time points *t* and *t*^*′*^, respectively. The model’s emphasis on maintaining similarity in gene representations across consecutive time points is determined by the scaling factor *α*, a hyperparameter we introduce to control the importance of temporal consistency within the graph. Additionally, *e*_*u,t*_ and *e*_*u,t*_*′* are the embeddings of node *u* at time points *t* and *t*^*′*^.

To illustrate how gene embeddings evolve over time, Figure 2 shows the trajectories of Node2vec embeddings for four representative genes of mRNA (VIM, Shisa3, EGFR, and Hist1h2ab) and four representative genes of miRNA (MYB, Zfp800, Nr6a1, and Hnrnpu) projected onto a 2D space using t-SNE [58]. With labels showing the historical order, each colored line represents a gene and demonstrates how its embedding changes over time and how the temporal Node2vec model captures patterns of similarity and differences in the dynamic behaviors of each gene. We combine the gene interaction graph and time-series gene expression data to create the temporal graph embeddings, as detailed in Algorithm 1.

**Figure 2.**
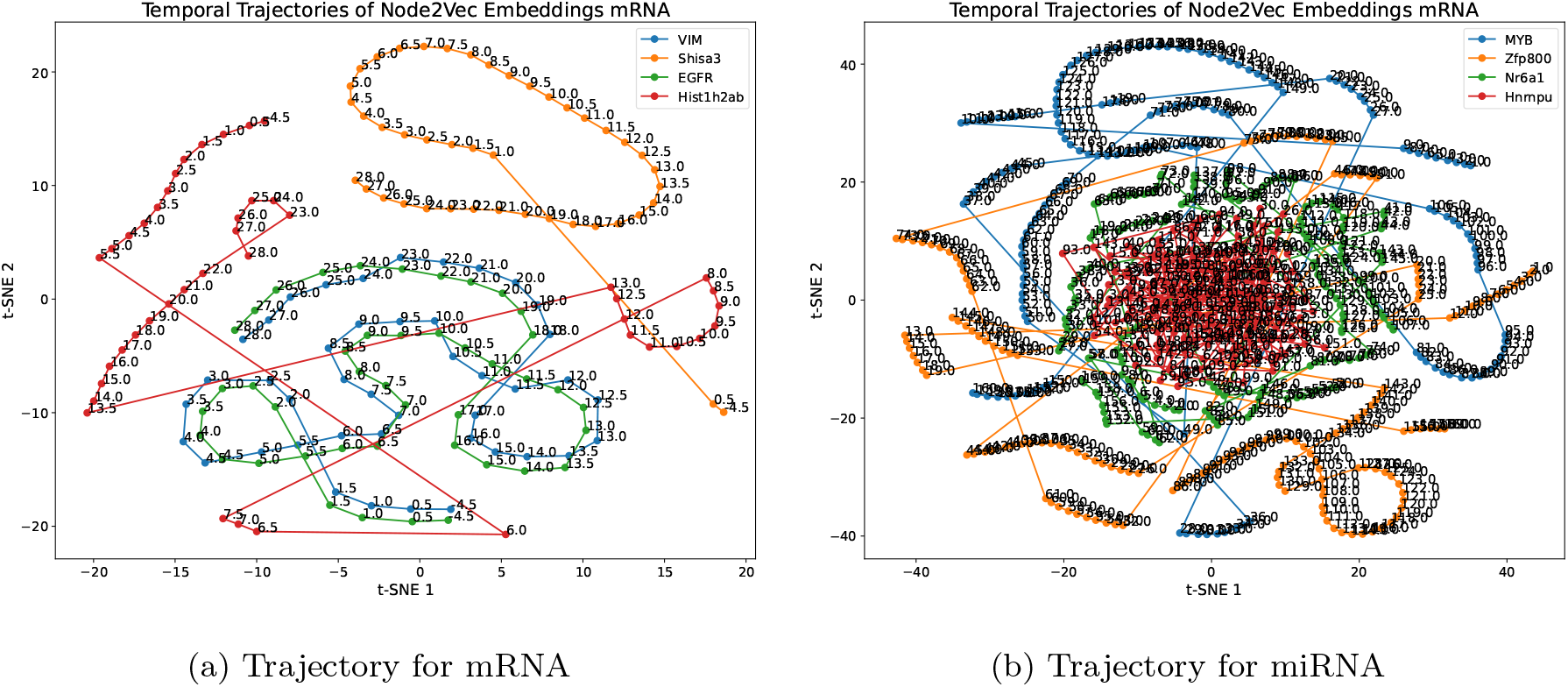
Temporal trajectories of Node2vec embeddings for selected genes, projected onto 2D space using t-SNE. The plots show four representative genes for mRNA (VIM, Shisa3, EGFR, Hist1h2ab) and four for miRNA (MYB, Zfp800, Nr6a1, Hnrnpu), illustrating how their learned Node2vec representations evolve over time.

#### Algorithm 1

Temporal Node2vec

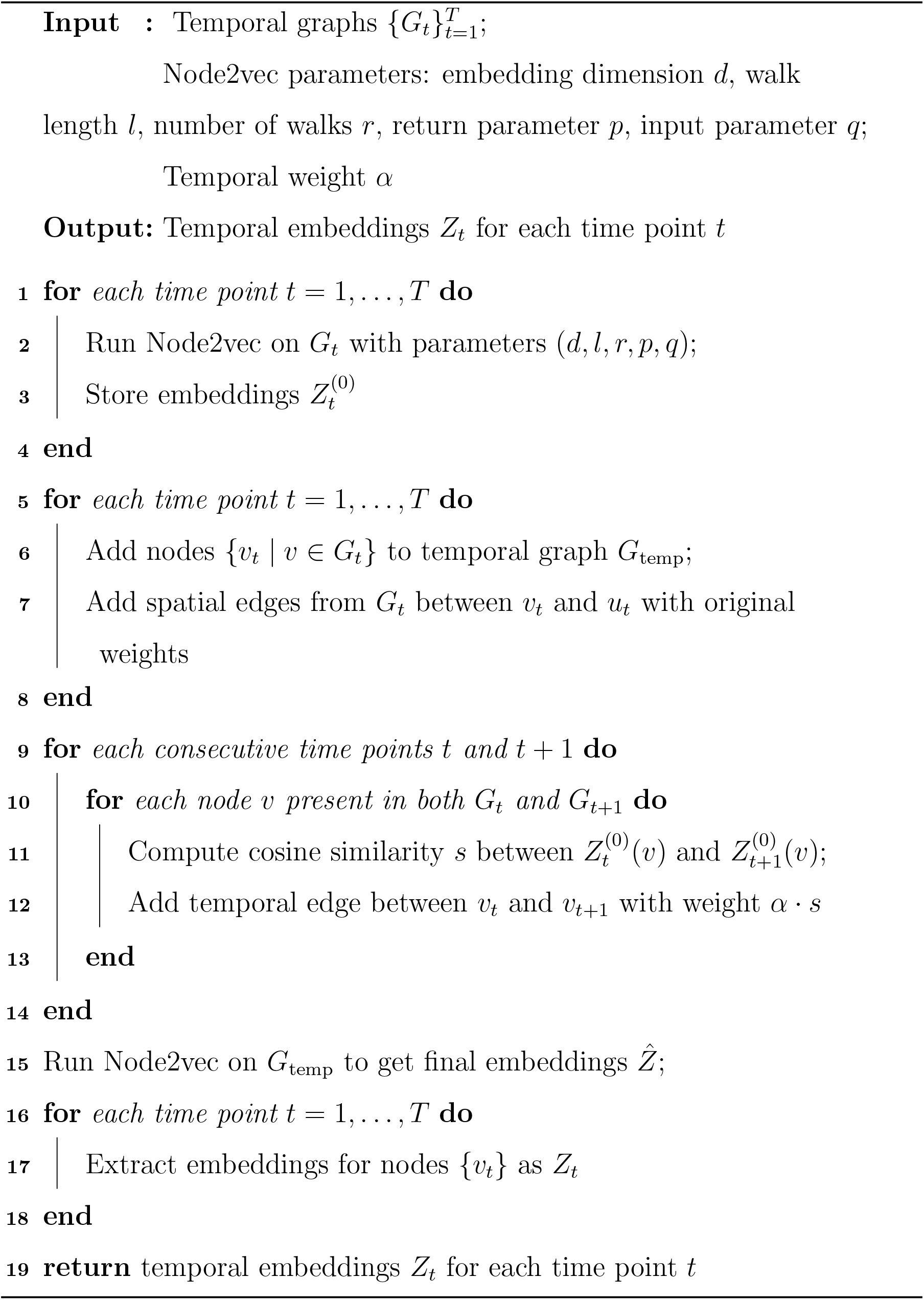

### 2.5. Towards STEPmr and STEPmi Architectures

We designed two distinct deep learning architectures to describe the temporal and structural dynamics of gene expression data, one architecture for mRNA expression prediction and the other for miRNA, since their target genes, interaction patterns, and the number of time points in datasets differ significantly. Both architectures are based on Spatio-temporal Graph Convolutional Networks (STGCNs) [59] with additional layers customized to each RNA type’s characteristics.

A 4D tensor of form ℝ^B×d×T ×N^ serves as the model’s input, where *B* is the batch size, *d* is the dimension of the Node2vec embedding vector for each node, *T* is the number of temporal steps, and *N* is the number of nodes (genes). A *d*-dimensional Node2vec embedding vector (see Section 2.4) is used to represent each node at each time step. The embeddings act as the input channels *C* = *d*, replacing raw expression values with structurally informed biological features. We use different values of *d* and *T* for miRNA and mRNA to represent their unique structural and temporal complexities.

### 2.5.1. Spatial-Temporal Graph Convolution Blocks (ST-Blocks)

At the core of the model is a sequence of Spatial Temporal Graph Convolution Block layers, each designed to comprehend both temporal and structural dependencies represented in the expression data.

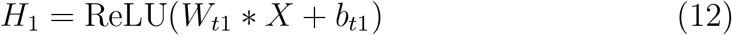

where ∗ denotes the temporal convolution operation with kernel size *K*_*t*_, and *W*_*t*1_ is the temporal kernel. This is followed by a graph convolution using Chebyshev polynomials of order *K*_*s*_, which encodes the structural dependencies among the nodes. The graph convolution is expressed with:

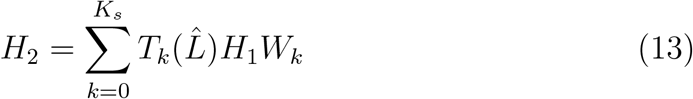

where 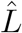 denotes the scaled graph Laplacian, *T*_*k*_ represents the Chebyshev polynomial of order *k* and *W*_*k*_ is the trainable weight matrix linked to the *k*-th polynomial term.

There is also a second temporal convolutional layer for both miRNA and mRNA after the graph convolutional layer with:

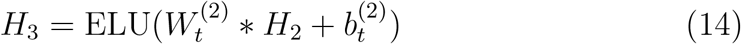

Here, ∗ denotes the temporal convolution. 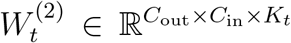 is the *temporal convolution kernel* for the second temporal layer, where *C*_in_ is the number of input channels as embedding dimension of Node2vec, *C*_out_ is the number of output channels, *K*_*t*_ is the temporal kernel size and 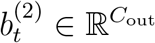 is the bias term.

#### 2.5.2. STEPmi (miRNA Specific Architecture)

After the spatio-temporal graph convolution blocks, the miRNA model (STEPmi) combines a multi-head self-attention module and a 4-layer bidirectional LSTM to enhance temporal and structural representations. The original features are integrated with the LSTM output using a residual connection and layer normalization given temporal input sequences 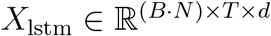, which represents the temporal sequences:

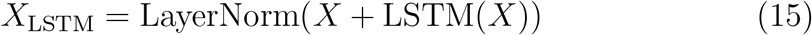

Long-range dependencies and context are captured by a 4-head selfattention layer, which helps to improve the model of dynamic gene interactions and overcome over-smoothing.

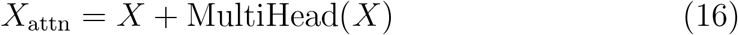

The general architecture of STEPmi is illustrated in Figure 3.

**Figure 3.**
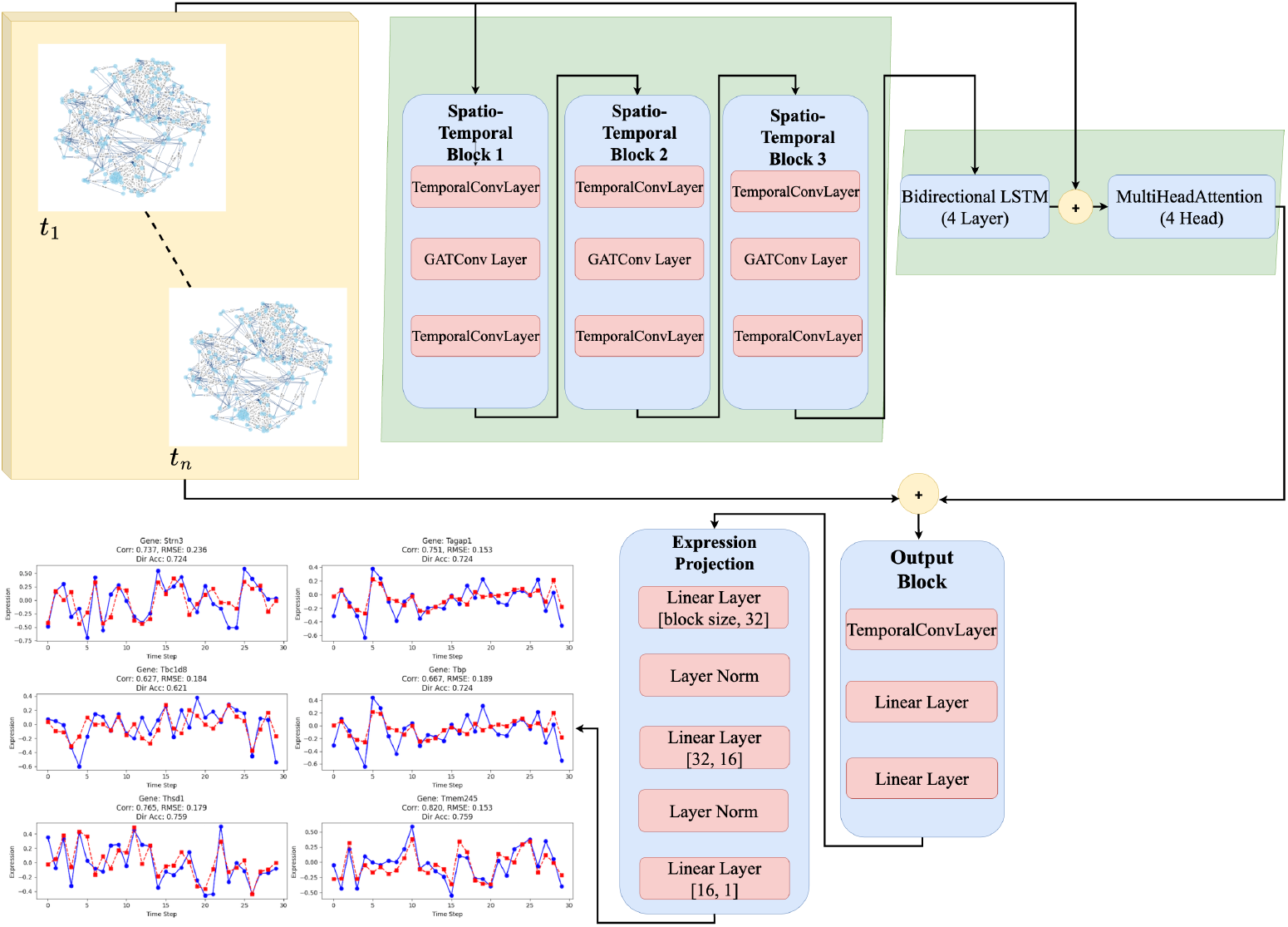
The architecture of STEPmi includes multi-head self-attention with four heads to capture complex and long-term interactions in gene expression, a 4-layer bidirectional LSTM for temporal enhancement, and spatio-temporal graph convolution blocks.

#### 2.5.3. STEPmr (mRNA Specific Architecture)

We modified the STGCN for mRNA architecture to include an attention mechanism based on gene connectivity values. Compared to miRNA, mRNA data is sparser and more prone to overfitting because it contains fewer time points (43) and target genes (49). So, STEPmr has a lightweight design appropriate for the limited mRNA dataset. mRNAs have more stable transcriptional behavior with a narrower regulatory scope than miRNAs, which influence several targets and exhibit dynamic, context-specific expression [60].

At each time step *t*, the model calculates an attention weight for every gene node *n* following the spatio-temporal convolutional blocks. This is accomplished by passing the node feature through a neural network. The result is a score *a*_*t,n*_ that indicates how important node *n* is at time *t*.

This attention score is modified using known gene-gene connection values *c*_*n*_ to account for previous biological knowledge. The final attention coefficient is obtained by scaling the influence of the connection with a learnable parameter, *γ*, with Equation 17:

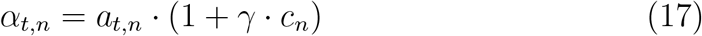

where *a*_*t,n*_ is the attention score for node *n* at time *t* and *c*_*n*_ is the connectivity weight. Equation 18 illustrates how a residual connection is used to calculate the final attention-modulated output in order to preserve the original signal and stabilize training.

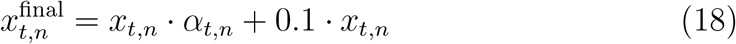

The overall architecture of the STEPmr architecture is illustrated in Figure 4.

**Figure 4.**
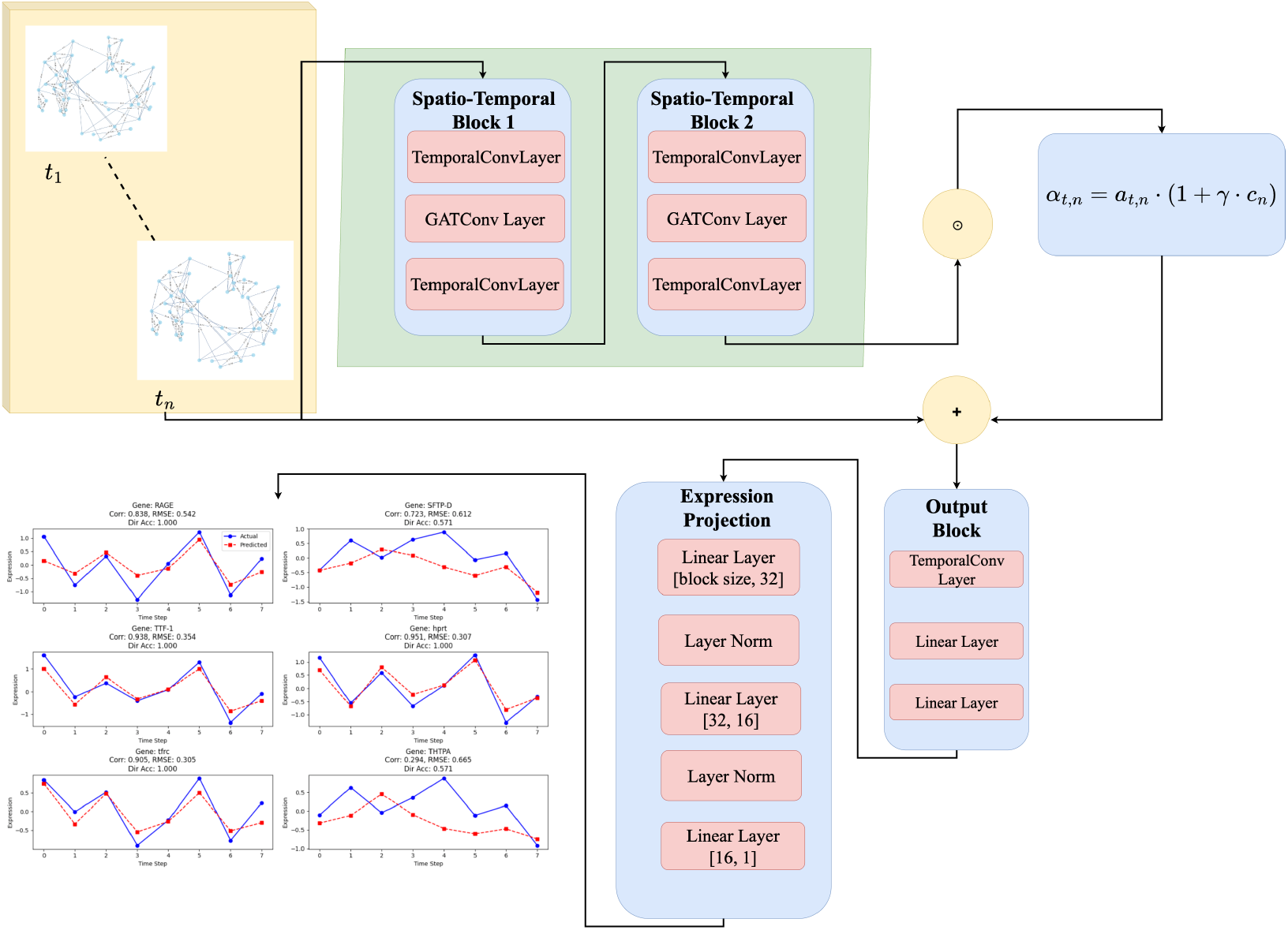
Architecture of STEPmr. STEPmr processes sparse temporal gene expression data using spatio-temporal graph convolutional layers, followed by a gene-aware attention. Gene-aware attention integrates biological gene-gene connectivity to clarify attention scores, which are used to generate gene expression predictions.

#### 2.5.4. Temporal-Aware Loss Functions

We designed a specific loss function beyond the conventional Mean Squared Error (MSE) to better reflect temporal dynamics in gene expression. MSE loss converges rapidly on large-scale data, and it is unable to identify temporal trends/directional changes, which are crucial in biological signals that change over time. Across both miRNA and mRNA datasets, our final loss is shown in Equation 19, which enables more biologically relevant predictions by combining MSE with additional terms that take trend alignment, change direction, and temporal coherence into account.

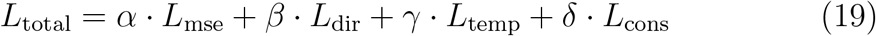

*L*_mse_ is mean squared error loss, *L*_dir_ represents the directional correction loss, *L*_temp_ is temporal trend correlation loss, and *L*_cons_ is the prediction consistency loss. Moreover, *α, β, γ, δ* are weighting coefficients for these losses.

##### MSE Loss

This is the primary term that penalizes differences between the ground truth expression values *Y* and the expected *Ŷ*:

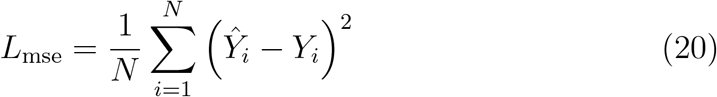

##### Directional Correction Loss

By penalizing inaccurate directionality, even when values are close, directional loss helps the model in learning temporal trends in gene expression. In order to penalize deviations from the expected trend, cosine similarity converts directional changes between time points into a loss by subtracting them from 1. The directional loss is given by:

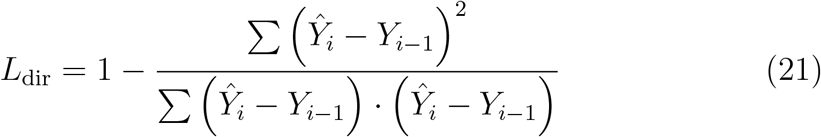

where *Ŷ*_*i*_ represents the predicted gene expression at time *t*_*i*_, and *Y*_*i*−1_ is the observed gene expression at the previous time step.

##### Temporal Trend Correlation Loss

We incorporate a Pearson correlation loss, which quantifies the degree to which the projected trajectory follows the real sequence, in order to match predicted gene expression with actual temporal patterns. This ensures consistent modeling of both individual and interacting gene behaviors. Temporal trend correlation loss calculated as:

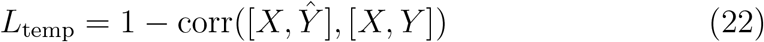

Here, [*X, Ŷ*] and [*X, Y* ] represent the concatenation of the input sequence *X* with the predicted *Ŷ* and true outputs *Y* .

##### Prediction Consistency Loss

Prediction consistency loss penalizes significant deviations from the previous expression value, ensuring that the model’s outputs remain consistent over time, particularly during short periods. This part is defined as:

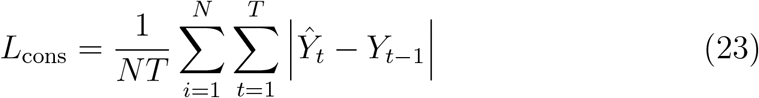

where *Ŷ*_*t*_ is the predicted gene expression at time step *t, Y*_*t*−1_ is the true gene expression at the previous time step *t* − 1, *N* is the total number of genes in the dataset and *T* is the total number of time points.

## 3. Results

### 3.1. Predictive Performance Metrics

We use two common correlation metrics, Pearson correlation and Spearman rank correlation, to evaluate our model’s temporal prediction performance on gene expression datasets. Both linear and monotonic relationships can be captured by these metrics, which enable us to measure the consistency of predicted and target expression levels.

Pearson correlation coefficient measures the linear relationship between the predicted gene expression values Ŷ and the corresponding ground truth values *Y* . Pearson correlation is calculated as follows:

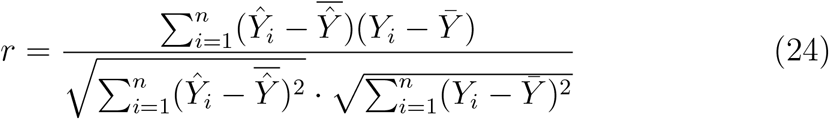

where *Ŷ*_*i*_ is the predicted expression value, *Y*_*i*_ is the true expression value at index *i*, and 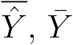 are respective means of ground truth predicted values. A high Pearson correlation suggests that the model successfully reflects the overall linear pattern of expression levels over time.

A non-parametric rank correlation, the Spearman correlation coefficient, evaluates how closely the relative ordering of expected gene expression values corresponds to the actual data. Spearman captures whether expression patterns and ranks are maintained over time, even if real values vary, in contrast to Pearson correlation, which quantifies linear relationships. Spearman correlation coefficient *ρ* is defined as:

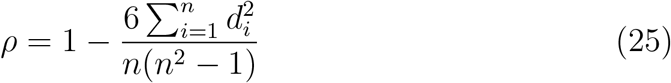

where *d*_*i*_ is the difference between the ranks of the predicted and true values for the *i*-th observation, and *n* is the total number of observations, where observations are expression values. In the context of temporal gene expression, a higher Spearman correlation indicates that the model preserves the ordered sequence of expression values, thus reflecting inherent coordinated and interaction relations among genes.

We also use MAE (Mean Absolute Error) and MSE (Mean Squared Error) as common regression metrics to assess the performance. Let *Y*_*t*_ be the true expression at time *t*, whereas *Ŷ*_*t*_ be the predicted expression at the same time step. They are defined as:

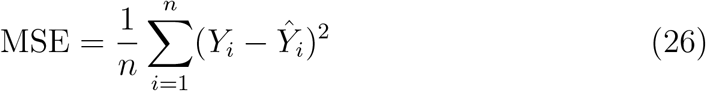

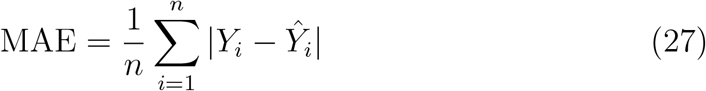

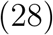

where lower scores mean better performance for both metrics.

### 3.2. Expression Predictability

We evaluate the expression prediction performance of STEPmi, by comparing it with a number of baseline models in terms of Mean Pearson and Mean Spearman correlations as shown in Table 1. For baseline approaches, if the baseline approach could not consider interactions, we tested them without taking interactions into account. Among the baselines, the Naive Baseline model uses only the expression value at the previous time point. Among the remaining baselines, Autoformer [61] is a highly used transformer-based technique for time series prediction. DCRNN (Diffusion Convolutional Recurrent Neural Network) [62] is a deep learning model for spatio-temporal forecasting on graph-structured data, especially designed for problems like traffic flow prediction. The remaining STGCN [59] and ASTGCN [51] are also spatio-temporal GNN-based approaches, mainly applied to multivariate time series prediction tasks other than biological tasks such as gene expression. According to the table, STEPmi achieves the best performance with 0.93 Pearson/Spearman correlations, respectively, outperforming the second-best performing approach STGCN by more than 15% in both metrics. Even though Autoformer is a transformer-based approach, its performance is quite low compared to STEPmi since it cannot take the interactions into account, which is one of the key factors in our performance. Two spatio-temporal GNN-based approaches, STGCN and ASTGCN, perform better than the remaining baselines since they can take the spatio-temporal nature of the data.

**Table 1:** Performance comparison with baseline models in terms of Mean Pearson and Mean Spearman correlations while predicting miRNA expression. The best performance is shown in bold.

| Model | Mean Pearson<br>(per gene over time) | Mean Spearman<br>(per gene over time) |
| --- | --- | --- |
| **STEPmi (Ours)** | **0.93** | **0.93** |
| Naive Baseline | 0.12 | 0.16 |
| Autoformer | 0.31 | 0.28 |
| DCRNN | 0.26 | 0.22 |
| STGCN | 0.80 | 0.80 |
| ASTGCN | 0.76 | 0.75 |

Similar to our approach for miRNA target genes, we also evaluated the performance of our approach STEPmr, by comparing it with the baseline approaches. The results are presented in Table 2. In general, the mRNA dataset had a relatively small number of target genes and time points, making baseline models more prone to overfitting. According to the table, STEPmr achieves the best performance with 0.77/0.74 Pearson/Spearman correlations, respectively, outperforming the second-best performing approach STGCN by almost more than 20% in both metrics. Similar to the results for STEPmi, Autoformer also performs relatively low compared since it cannot take the interactions into account. Two spatio-temporal GNN-based approaches, STGCN and ASTGCN, also perform better than the remaining baselines since they can take the spatio-temporal nature of the data.

**Table 2:** Performance comparison with baseline models in terms of Mean Pearson and Mean Spearman correlations while predicting mRNA expression. The best performance is shown in bold.

| Model | Mean Pearson<br>(per gene over time) | Mean Spearman<br>(per gene over time) |
| --- | --- | --- |
| **STEPmr (Ours)** | **0.77** | **0.74** |
| Naive Baseline | -0.11 | -0.09 |
| DCRNN | 0.23 | 0.20 |
| Autoformer | 0.38 | 0.38 |
| STGCN | 0.51 | 0.51 |
| ASTGCN | 0.58 | 0.52 |

The actual and predicted expression trajectories for the five genes with the highest mean Spearman/Pearson correlation among the target genes in the miRNA and mRNA datasets are shown in Figures 5-6, respectively. In this case, after training the expression miRNA dataset with STEPmi, we evaluated the model on 32 unseen time points. For every gene, we provide predicted trajectories, Pearson correlation, and RMSE. The results show how temporal loss and STEPmi’s design work together to accurately replicate gene expression dynamics and preserve actual temporal patterns. Although miRNA and mRNA have different regulation mechanisms, which result in different absolute expression values, the five genes with the highest performance in each group show consistent temporal patterns. These genes exhibit strong alignment in the way their expression changes over time, although essentially differing in magnitude and underlying biological properties.

**Figure 5.**
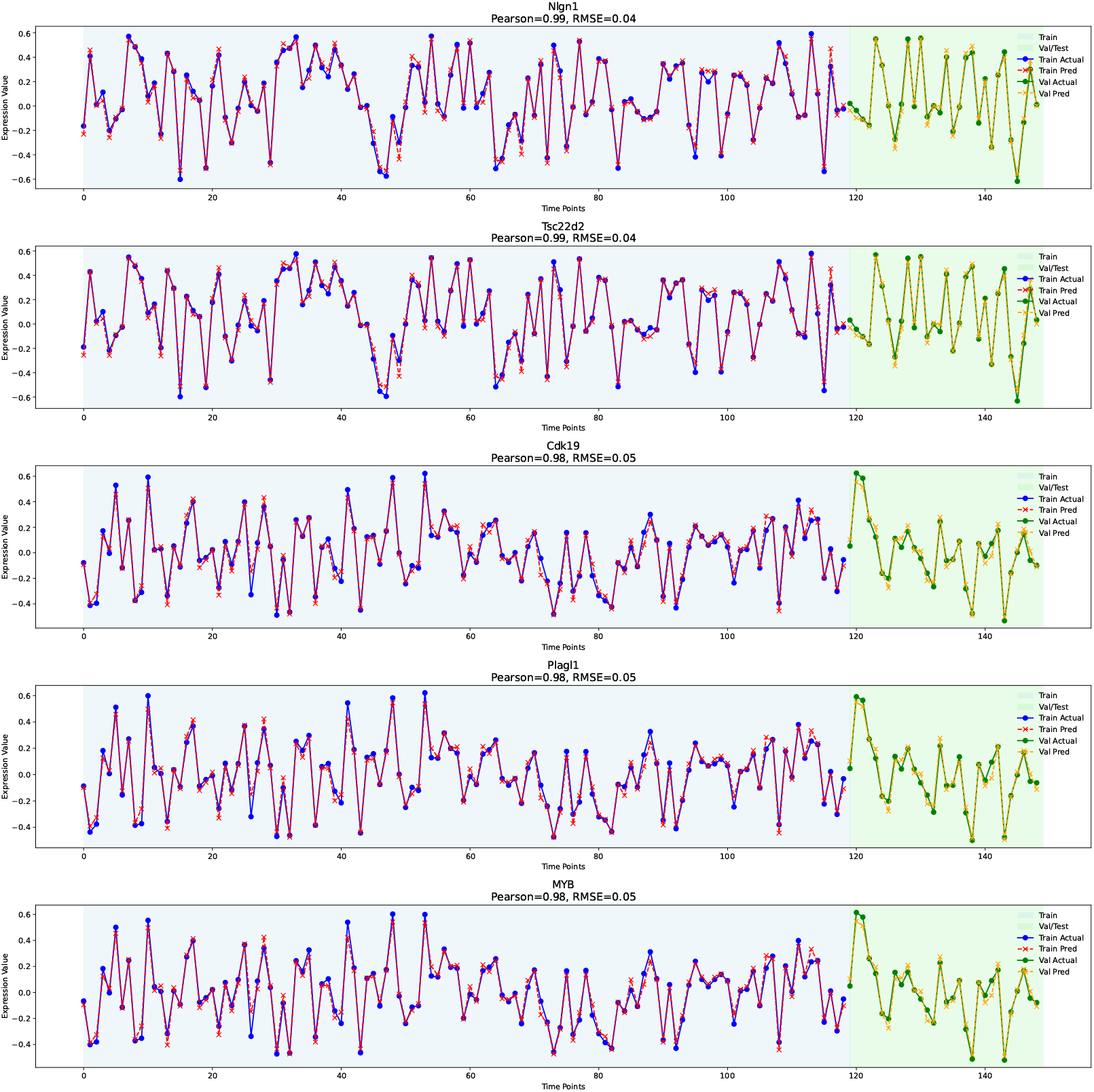
Predicted and actual expression trajectories for the top five performed miRNA genes on the train/test set. Each subplot presents the ground truth and predicted values over 32 unseen time points. Pearson correlation and RMSE are reported for each gene.

**Figure 6.**
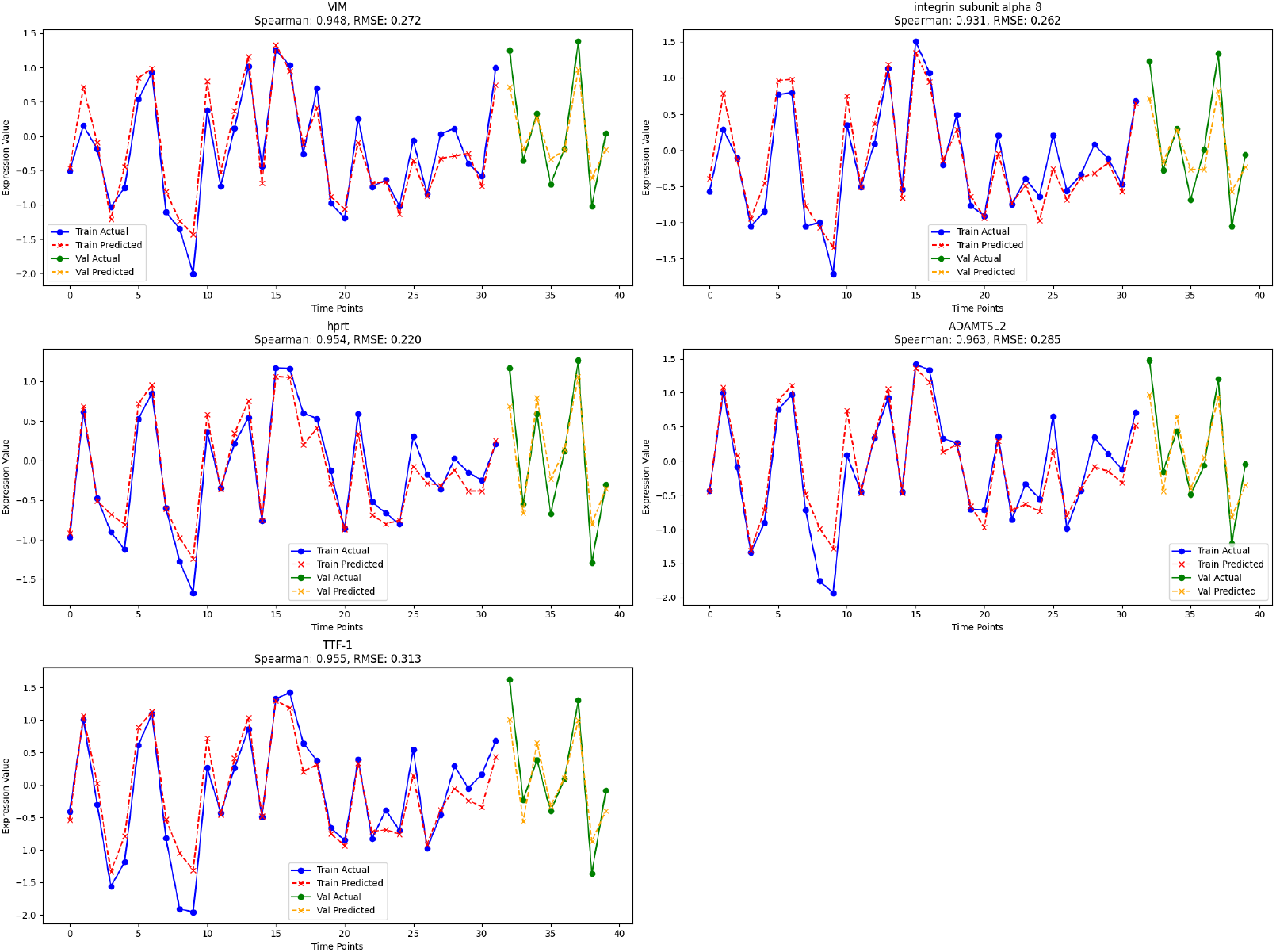
Predicted and actual expression trajectories for the top five performed mRNA genes on the train/test set. Each subplot presents the ground truth and predicted values over 8 unseen time points. Pearson correlation and RMSE are reported for each gene.

To investigate the relationships as well as co-expression patterns among the best-performing genes, we additionally compute the Pearson correlation values between the top 8 best-performing genes of miRNA and mRNA, as shown in Figures 7-8 respectively. In general for both datasets, we find these top-performing genes to be highly correlated across time points, which suggests their shared role in multiple cellular processes and regulatory pathways [63].

**Figure 7.**
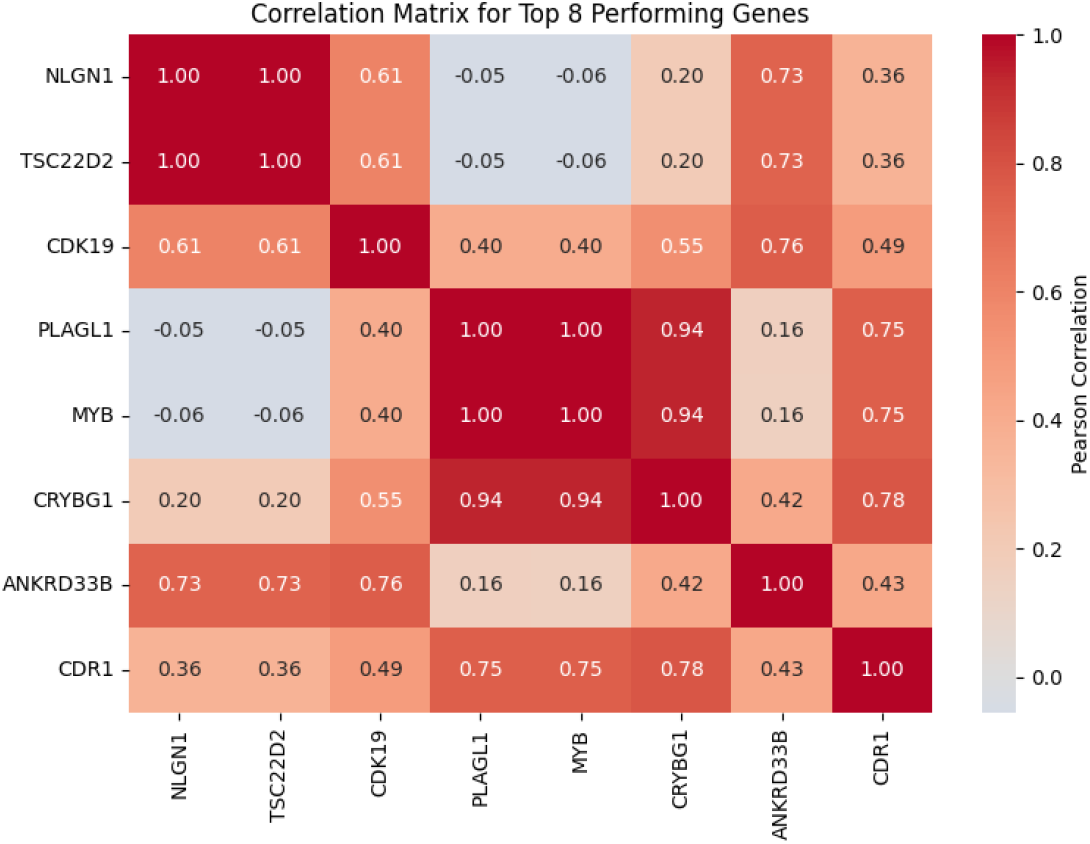
Correlation matrix showing the pairwise Pearson correlation coefficients between the top 8 highest performing genes for miRNA.

**Figure 8.**
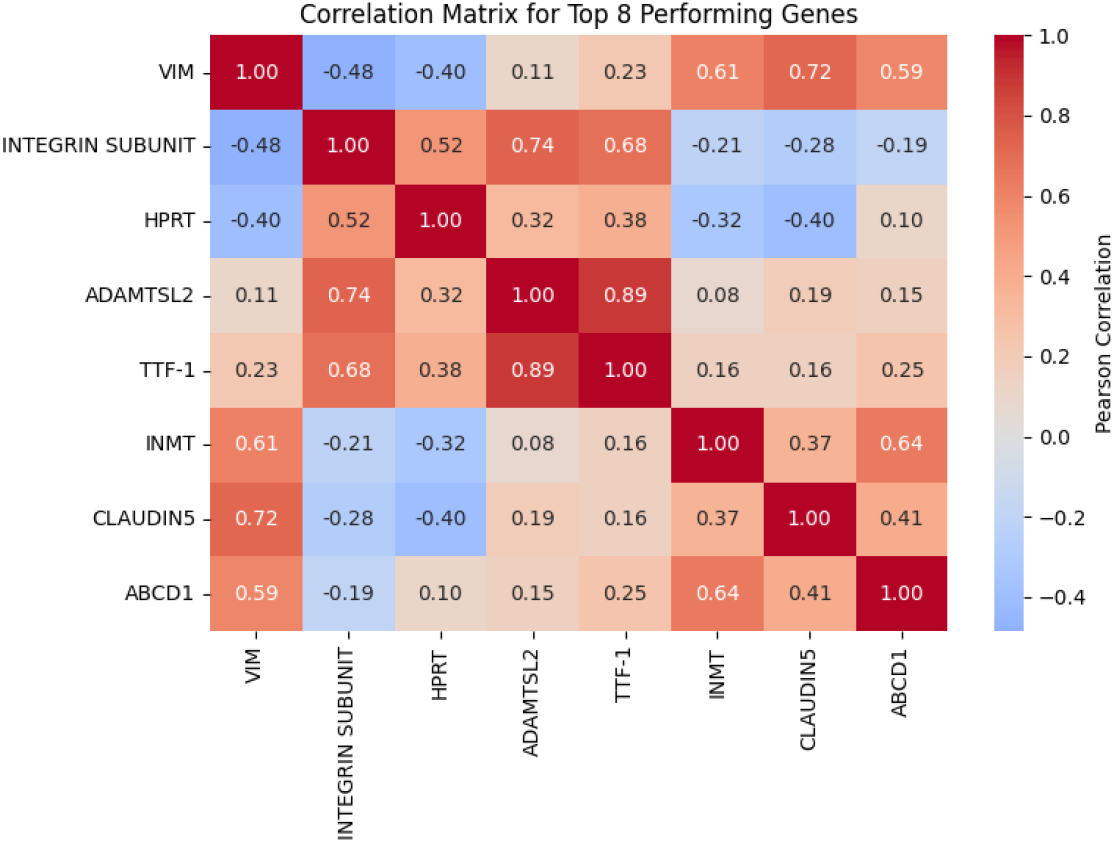
Correlation matrix showing the pairwise Pearson correlation coefficients between the top 8 highest performing genes for mRNA.

### 3.3. Ablation Studies

We analyze the impact of temporal-aware loss on the performance of STEPmi and STEPmr. As seen in Table 3, we obtain better performance by using a temporal-aware loss in both approaches since adding temporal constraints on time series prediction helps improve the performance via regularization-like impact. When we use only MSE loss, it is unable to identify temporal trends/directional changes, which are crucial in biological signals that change over time. Temporal-aware loss is more effective in STEPmi, whereas it only slightly increases the performance in STEPmr. Such a performance difference between the two approaches can be attributed to different temporal and spatial characteristics of the datasets. On the other hand, Table 4 analyzes the prediction performance by using static or temporal Node2vec. In both datasets, temporal NodeVec is significantly more effective than static/traditional Node2vec, which shows the importance of the temporal embedding component in our methods.

**Table 3:** The impact of temporal-aware loss in both STEPmi and STEPmr, by comparing its performance with MSE case.

| Model | Mean Pearson<br>(per gene over time) | Mean Spearman<br>(per gene over time) |
| --- | --- | --- |
| **STEPmi with temporal-aware loss** | **0.93** | **0.93** |
| STEPmi with MSE | 0.86 | 0.84 |
| **STEPmr with temporal-aware loss** | **0.77** | **0.74** |
| STEPmr with MSE | 0.76 | 0.72 |

**Table 4:** The impact of using static vs temporal Node2vec in STEPmi and STEPmr performance.

| Model | Node2vec type | Mean Pearson | Mean Spearman |
| --- | --- | --- | --- |
|  |  | (per gene over time) | (per gene over time) |
| STEPmi | Static | 0.67 | 0.60 |
| **STEPmi** | **Temporal** | **0.93** | **0.93** |
| STEPmr | Static | 0.62 | 0.50 |
| **STEPmr** | **Temporal** | **0.77** | **0.72** |

As another ablation study, we focused on the impact of individual components within both STEPmi and STEPmr. According to Table 5, ST blocks are important for both methods. Moreover, bidirectional LSTM is also important in STEPmr performance. Removing the individual model components decreases the spatio-temporal expression prediction performance in terms of multiple metrics, including MSE and MAE as well.

**Table 5:** Performance comparison of STEPmr and STEPmi architectures under different configurations: full model vs. models with reduced ST blocks or without connectivity attention, BiLSTM, or multi-head attention.

| Model | Pearson | Spearman | MSE | MAE |
| --- | --- | --- | --- | --- |
| **STEPmr** | **0.77** | **0.74** | **0.21** | **0.37** |
| STEPmr one ST block | 0.69 | 0.65 | 0.25 | 0.38 |
| STEPmr w/o Connectivity Attention | 0.68 | 0.62 | 0.26 | 0.39 |
| **STEPmi** | **0.93** | **0.93** | **0.01** | **0.06** |
| STEPmi one ST block | 0.84 | 0.84 | 0.04 | 0.09 |
| STEPmi two ST block | 0.85 | 0.84 | 0.04 | 0.08 |
| STEPmi w/o BiLSTM | 0.90 | 0.90 | 0.03 | 0.08 |
| STEPmi w/o Multi Head Attention | 0.89 | 0.90 | 0.02 | 0.08 |

### 3.4. The Impact of Hyperparameters

For both STEPmi and STEPmr, we evaluated different dimensions for Node2vec embeddings in order to assess how embedding dimensionality affects model performance. These dimensions change the input size of the ST-Blocks as well as the structure of temporal node representations since the embeddings are integrated into the interaction graph. We explored 64, 128, 256, and 512 embedding dimensions for STEPmi, and 16, 32, and 64 dimensions for STEPmr. Only the input dimensionality of the ST-Block was changed for a fair comparison, while all other hyperparameters remained unchanged. For STEPmi, the model that used 256-dimensional Node2vec embeddings gives the highest mean Pearson and Spearman correlation, as shown in Table 6. On the other hand, the unique characteristics of the mRNA dataset necessitated a distinct range of embedding dimensions compared to those used for the miRNA dataset. In particular, smaller embedding sizes were taken into consideration since higher-dimensional embeddings (e.g., 128 or above) would be inappropriate for this dataset as they would greatly exceed the total number of available time points and target genes. The results of embedding dimensionality for mRNA are summarized in Table 7.

**Table 6:** The impact of Node2vec embedding dimensionality on the performance in STEPmi.

| Embedding Dimension | Mean Pearson<br>(per gene over time) | Mean Spearman<br>(per gene over time) |
| --- | --- | --- |
| 64 dimensional Node2vec | 0.83 | 0.83 |
| 128 dimensional Node2vec | 0.85 | 0.83 |
| **256 dimensional Node2vec** | **0.93** | **0.93** |
| 512 dimensional Node2vec | 0.73 | 0.71 |

**Table 7:** The impact of Node2vec embedding dimensionality on the performance in STEPmr.

| Embedding Dimension | Mean Pearson<br>(per gene over time) | Mean Spearman<br>(per gene over time) |
| --- | --- | --- |
| 16 dimensional Node2vec | 0.45 | 0.40 |
| **32 dimensional Node2vec** | **0.77** | **0.74** |
| 64 dimensional Node2vec | 0.48 | 0.45 |

The development of graph-based embeddings is primarily dependent on the Node2vec parameters rather than the expression data itself, despite miRNA and mRNA expression levels exhibiting different temporal patterns. The influence of various hyperparameter configurations on Node2vec-based embedding creation was examined by maintaining a constant walk length and number of walks while varying the return parameter *p* and the in–out parameter *q*. These parameters regulate the random walk behavior: *q* biases the walk toward exploring new or distant nodes, while *p* controls the likelihood of returning to a node in the walk. We tested scenarios with different parameter relations, such as *p > q, q > p*, walk length *>* number of walks, and number of walks *>* walk length. From this parameter search, the best performance was achieved with the configuration *p* = 1.0, *q* = 0.5, walk length = 25, and number of walks = 75, resulting in a Mean Pearson correlation of 0.93 and a Mean Spearman correlation of 0.93 per gene over time can be seen from Table 8. For mRNA dataset, the highest performance values were obtained with the embedding parameters: *p* = 1.0, *q* = 1.0, walk length = 25, and number of walks = 75. The correlation values obtained with different hyperparameters for the mRNA dataset can be seen in Table 9. In essence, these Node2vec parameters specify the embedding training approach, showing that they successfully capture the structural and temporal associations that are needed for precise expression prediction in spite of biological variations.

**Table 8:** The performance of STEPmi over various Node2vec hyperparameter combinations.

| Node2vec Parameters | Mean Pearson<br>(per gene over time) | Mean Spearman<br>(per gene over time) |
| --- | --- | --- |
| p = 0.5, q = 0.5, walk_len = 25, num_walks = 75 | 0.90 | 0.88 |
| p = 1.5, q = 1.5, walk_len = 25, num_walks = 75 | 0.93 | 0.92 |
| p = 2.0, q = 1.0, walk_len = 25, num_walks = 75 | 0.93 | 0.90 |
| **p = 1.0, q = 0.5, walk_len = 25, num_walks = 75** | **0.93** | **0.93** |
| p = 1.0, q = 2.0, walk_len = 25, num_walks = 75 | 0.76 | 0.74 |
| p = 1.0, q = 1.0, walk_len = 25, num_walks = 75 | 0.90 | 0.89 |
| p = 1.0, q = 1.0, walk_len = 20, num_walks = 90 | 0.90 | 0.89 |
| p = 1.0, q = 1.0, walk_len = 30, num_walks = 60 | 0.89 | 0.87 |
| p = 1.0, q = 1.0, walk_len = 30, num_walks = 90 | 0.85 | 0.84 |
| p = 1.0, q = 1.0, walk_len = 35, num_walks = 50 | 0.88 | 0.85 |
| p = 1.0, q = 1.0, walk_len = 80, num_walks = 10 | 0.86 | 0.84 |

**Table 9:** The performance of STEPmi over various Node2vec hyperparameter combinations.

| Node2vec Parameters | Mean Pearson<br>(per gene over time) | Mean Spearman<br>(per gene over time) |
| --- | --- | --- |
| **p = 1.0, q = 1.0, walk_len = 25, num_walks = 75** | **0.77** | **0.72** |
| p = 1.5, q = 1.5, walk_len = 25, num_walks = 75 | 0.45 | 0.43 |
| p = 2.0, q = 1.0, walk_len = 25, num_walks = 75 | 0.73 | 0.69 |
| p = 1.0, q = 0.5, walk_len = 25, num_walks = 75 | 0.56 | 0.55 |
| p = 1.0, q = 2.0, walk_len = 25, num_walks = 75 | 0.48 | 0.51 |
| p = 1.0, q = 1.0, walk_len = 20, num_walks = 90 | 0.55 | 0.45 |
| p = 1.0, q = 1.0, walk_len = 30, num_walks = 60 | 0.49 | 0.41 |
| p = 1.0, q = 1.0, walk_len = 30, num_walks = 90 | 0.71 | 0.68 |
| p = 1.0, q = 1.0, walk_len = 35, num_walks = 50 | 0.47 | 0.39 |
| p = 1.0, q = 1.0, walk_len = 80, num_walks = 10 | 0.33 | 0.28 |

### 3.5. GO Enrichment Analysis for Biological Interpretation

We used Enrichr API [64] to perform Gene Ontology (GO) enrichment analysis on the target genes of miRNA and mRNA independently, aiming to reveal the biological significance of our model’s predictions. We classified the miRNA and mRNA target genes based on several factors, such as their prediction performance, chromatin structure (A/B compartments), topologically associating domain (TAD) boundaries, and temporal expression patterns (upregulated, stable, or downregulated). For each category, we identified relevant GO molecular functions and biological processes. Additionally, we examined GO terms for the five genes with the highest and the lowest performance in order to investigate possible biological factors that could influence their prediction performance.

#### 3.5.1. Ontology-Based Interpretation of miRNA Target Gene Groups

To clarify the biological implications of our model, we ranked target genes based on their Pearson and Spearman correlations within the test set and selected the highest and lowest-performing ones. Since even the gene with the lowest performance showed an 86% correlation, demonstrating the model’s overall good performance, we concentrated on presenting miRNA target genes first for the biological results. Furthermore, the miRNA dataset provides adequate coverage to generate Gene Ontology terms that offer significant biological insights, given its total of 162 genes and 159 time points. These choices provide insight into which gene expression patterns the model captures accurately and which ones are still challenging to predict. Based on Gene Ontology (GO) terminology, we created separate tables for genes in different categories (including the highest-lowest correlated gene tables), describing their primary biological and molecular roles. Trends linked to strong or weak model performance were also identified by looking at known regulatory characteristics and temporal expression patterns. For the lowest-correlated genes, we also considered possible external influences that might have affected prediction quality.

The five best-performing miRNA target genes are shown in Table 10 to help examine the model’s performance in terms of gene-specific features. Tissue-specific expression patterns, fundamental molecular activities, expression behavior across time, regulatory characteristics, and important biological roles are all included in each entry. For instance, MYB, one of the highest-performing genes, belongs to strong TAD boundaries within the B compartment, which indicate structurally stable genomic regions, and it shows stable expression in several immune cell lineages [65]. Other high-performing genes, like NLGN1, TSC22D2, CDK19, and PLAGL1, also inhabit genomic areas that are suitable to reliable regulation and are involved in essential transcriptional and stress-response pathways [66, 67]. With consistent temporal and functional patterns, these observations show that the model can acquire significant regulatory and expression aspects from the miRNA dataset, precisely representing the biological behaviors of genes.

**Table 10:** Highest performing 5 genes for miRNA.

| Gene | Tissue-specificity | Primary Function | Key Biological Features |
| --- | --- | --- | --- |
| **NLGN1**<br>Expression Behaviour: Downregulated<br>Regulatory Features: Weak TAD boundaries, A compartment | macrophage bone marrow 2hr LPS, dorsal striatum | Synaptic cell adhesion, neuron-neuron signaling, immune response modulation | Regular participation in immune regulation/-plasticity of synaptic neurons, synchronized downregulation during inflammation |
| **TSC22D2**<br>Expression Behaviour: Downregulated<br>Regulatory Features: Strong and Weak TAD boundaries, A/B compartments | MEF, mast cells IgE+antigen 1hr | Transcriptional repressor, stress response modulation, differentiation | affects immune gene expression and differentiation processes, takes role in stress-responsive transcriptional networks |
| **CDK19**<br>Expression Behaviour: Upregulated<br>Regulatory Features: Strong TAD boundaries, A compartment | prostate | RNA polymerase II CTD phosphorylation, cyclin-dependent protein kinase activity, protein modification | Upregulated expression pattern linked to fundamental transcriptional machinery, essential function in RNA polymerase II regulation |
| **PLAGL1**<br>Expression Behaviour: Stable<br>Regulatory Features: Strong and Weak TAD boundaries, A/B compartments | pituitary, umbilical cord | DNA damage response with p53 signaling, cell cycle regulation, transcriptional activation/repression, apoptotic process | Stable expression linked to fundamental cellular processes, consistent role in cell cycle control |
| **MYB**<br>Expression Behaviour: Stable<br>Regulatory Features: Strong TAD boundaries, B compartment | Baf3, thymocyte DP CD4+CD8+, bone marrow | Transcriptional regulation of hematopoiesis, T-helper cell differentiation control, histone methylation (H3-K9/H3-K4) regulation | Consistent expression in immune cell lineages, regulated epigenetic modification patterns, stable transcriptional activity across developmental stages |

The five miRNA target genes with the lowest performance are shown in Table 11. These genes capture biologically significant associations despite having lower performance based on correlation than the top-performing group (lowest with %86 correlation), demonstrating the model’s ability to capture complex and context-dependent expression patterns. Tissue-specific expression, primary function, context-dependent expression modified by external stimuli, and possible causes of low performance are summarized in each entry. For example, membrane potential and synaptic signaling have an impact on the upregulated, activity-driven expression of DPP10 in neuronal tissues, which restricts predictability across a variety of tissues [68]. Likewise, the high context-specific regulation of DUSP23, PHLPP1, KLHDC8A, and KLF7 genes—including their reactions to stress, developmental cues, inflammation response, and signaling pathways—contributes to their lower performance [69, 70]. We can observe from the lowee performing gene analysis that the model achieves a certain amount of biological relevance but struggles to predict highly variable and stimulus-dependent expression values.

**Table 11:**
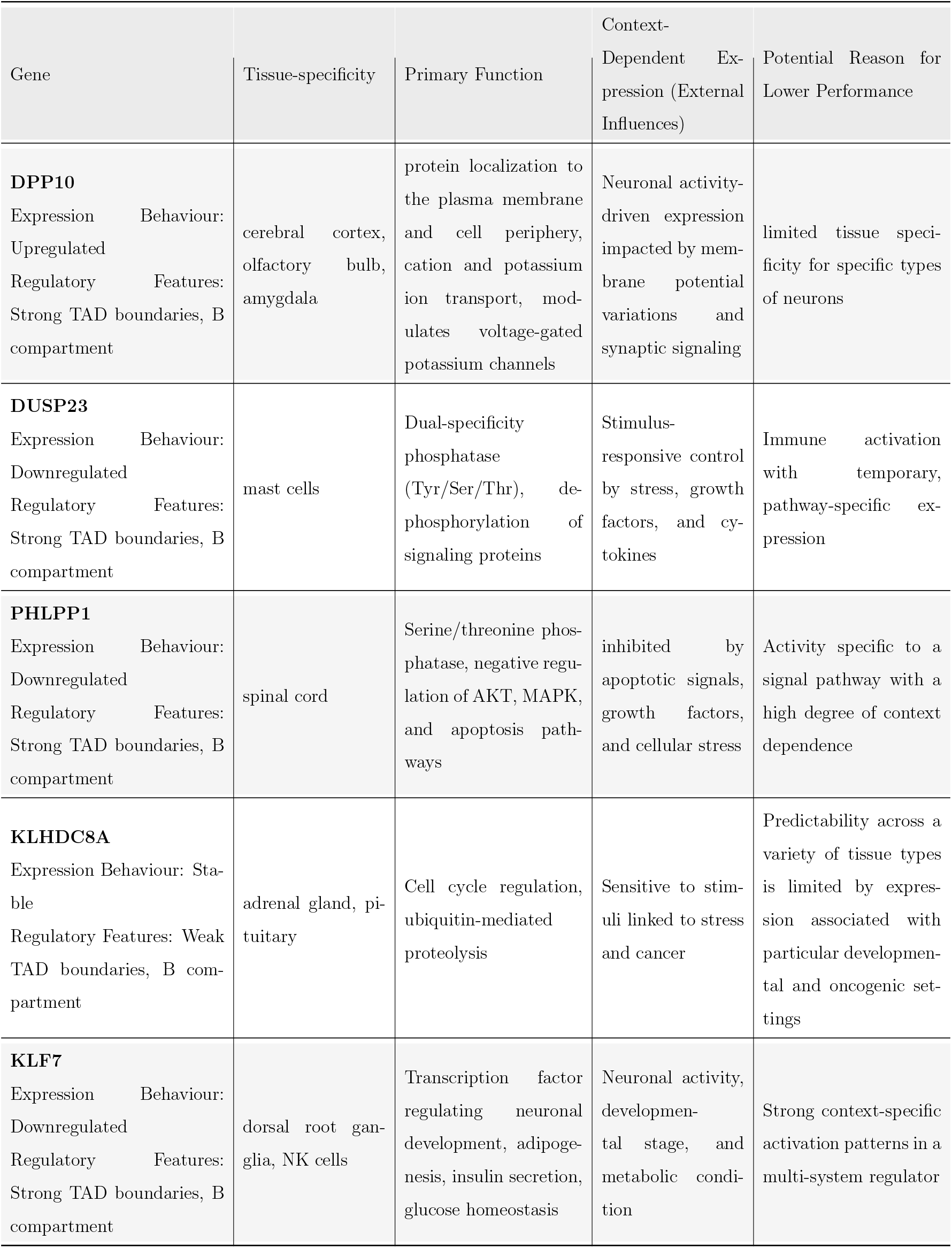
Lowest performing 5 genes for miRNA.

Examining the miRNA target genes with the highest and lowest performance provides a thorough understanding of the model’s behavior. Genes with the best performance typically show consistent or stable expression patterns across tissues and developmental stages, participate in essential biological functions (e.g., transcriptional regulation, RNA processing, structural maintenance), and are controlled within robust genomic areas. Lower performing genes, on the other hand, frequently exhibit extremely dynamic, context-dependent expression, participate in intricate or condition-specific processes like tissue remodeling, immune response, oxidative stress, or cellcycle–related signals. The outcomes indicate that the model, which offers both prediction accuracy and biological interpretability, not only detects significant trends in gene expression but also reflects the biological restrictions and variability found in the miRNA dataset.

We expand our biological interpretation by performing Gene Ontology enrichment analysis on the downregulated gene cluster to investigate relevant molecular functions and biological processes. The significance level of each pathway was determined based on the enrichment rank and statistical strength of the GO annotations. We concluded by highlighting key genes, which are the most representative or functionally central in each pathway, to support the inferred biological roles. For example, NLGN1 and KLF7 are key regulators of synapse assembly and neurotransmitter dynamics [71, 72], where those genes are included in neuronal development and synaptic function with high significance that can be seen in Table 12. All things considered, the downregulated gene cluster is filled with critical cellular and developmental pathways, especially those related to membrane organization, immunological regulation, and neural development. Pathways related to neurodevelopment and synapse function in particular show very high importance, indicating that these processes are active at the time points used for model training and evaluation.

**Table 12:**
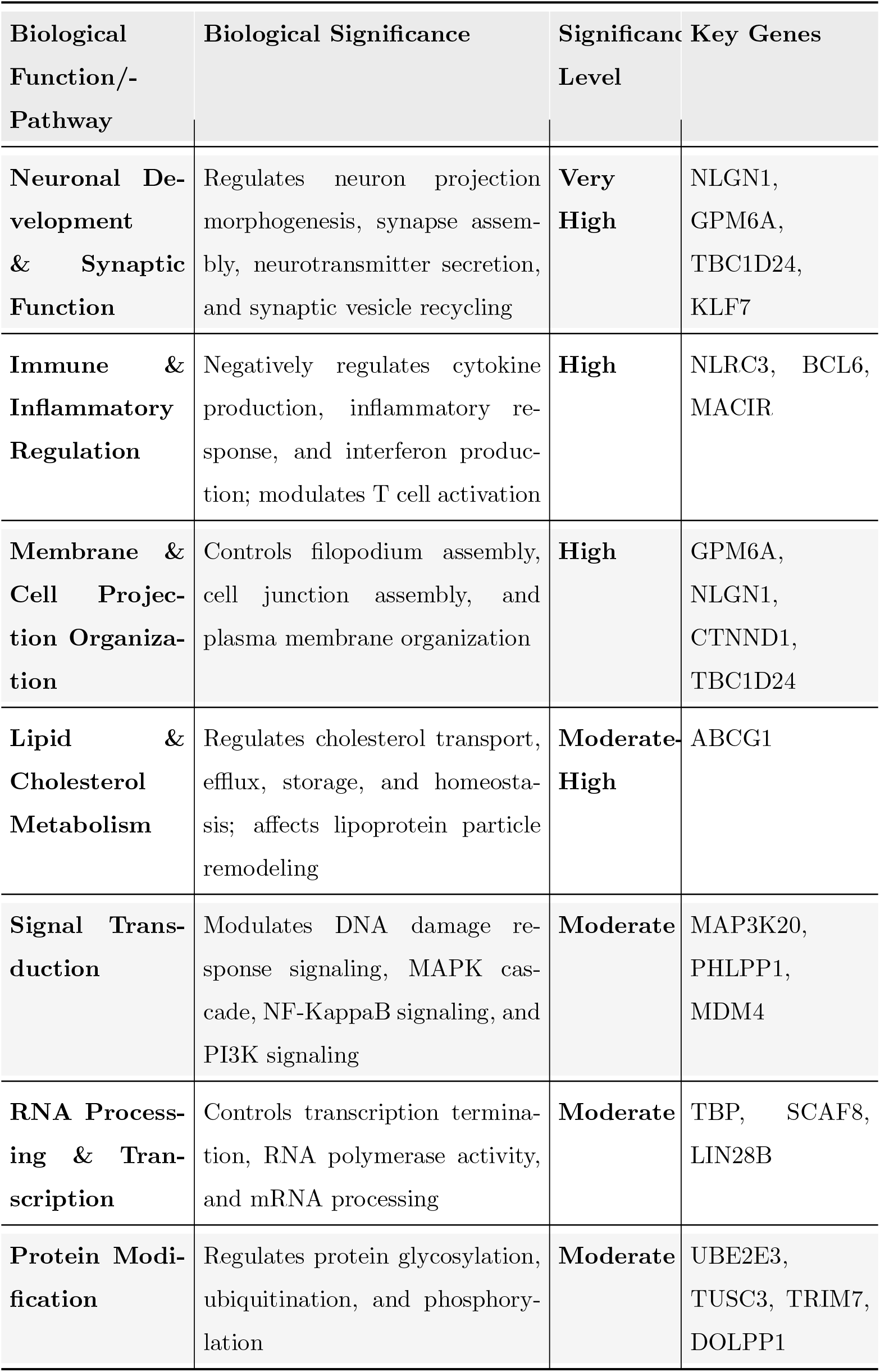
Biological Pathway Significance of Downregulated Genes.

The downregulated gene’s molecular function analysis indicates enrichment in categories essential for membrane dynamics, signal transduction, and transcriptional regulation. According to the molecular function Table 13, downregulated genes are substantially more common in roles related to lipid transport and RNA polymerase II machinery interaction, suggesting that essential processes like transcription initiation and lipid handling may be particularly affected. Further downregulation of vital regulatory, signaling, and transport pathways is indicated by the suppression of other activities, including transport, kinase and phosphatase activity, and membrane protein interactions. These findings imply that the downregulation affects fundamental molecular processes essential for communication and cellular maintenance. Stable and upregulated gene clusters were analyzed for miRNA enrichment using the same methodology, and the detailed results are provided in the Supplementary Material.

**Table 13:** Molecular Function Significance of Downregulated Genes.

| Molecular Function Category | Biological Significance | Significance Level | Key Genes |
| --- | --- | --- | --- |
| **RNA Polymerase II Machinery Interaction** | Involved in transcription machinery binding, C-terminal domain binding, and transcription initiation factor binding | **Very High** | TBP, SCAF8 |
| **Lipid and Cholesterol Transport** | Mediates phosphatidylcholine floppase activity, cholesterol transfer, and sterol transport | **High** | ABCG1 |
| **Immune Signaling Regulation** | Regulates phosphatidylinositol 3-kinase regulatory subunit binding and immune response pathways | **High** | NLRC3 |
| **Ion Transport** | Functions in magnesium ion transmembrane transport and metal ion transport | **Moderate-High** | TUSC3 |
| **Phosphatase Activity** | Exhibits protein tyrosine/serine/threonine phosphatase activity and phosphoric ester hydrolase activity | **Moderate** | DUSP23, PHLPP1 |
| **Kinase Activity** | Shows MAP kinase kinase activity and protein tyrosine kinase activity | **Moderate** | MAP3K20, ITK |
| **Membrane Protein Interaction** | Facilitates neurexin family protein binding and PDZ domain binding | **Moderate** | NLGN1 |
| **Nucleotide Binding** | Involves ADP binding, ATP binding, and purine ribonucleoside triphosphate binding | **Low-Moderate** | ABCG1, GNL1 |
| **DNA Binding** | Functions in sequence-specific DNA binding and transcription factor activity | **Low** | TBP, BCL6, FLI1, KLF7, SATB2 |

By comparing the molecular and functional characteristics of genes located in strong and weak TAD boundaries, we evaluated the biological significance of TAD boundary strength. Genes within strong TAD boundaries primarily participate in transcriptional regulation, immune system function, cell structure organization, and neuronal processes, indicating a compartmentalized regulatory role in cellular functions. On the other hand, genes that are enriched for RNA processing, neuronal development, transcriptional control, and signal transduction pathways are found within weak borders; this indicates a more dynamic and diverse regulatory context. Detailed categorizations of these functional enrichments can be found in Tables 14 and 15. For a closer examination, we highlighted the ‘Transcriptional Regulation’ entry in the strong TAD boundary gene table, detailing the unique functional and molecular attributes associated with these genes. These genes play a major role in RNA processing, mRNA stabilization, and miRNA regulation, which correspond to processes that are essential for maintaining cellular homeostasis and controlling gene expression. This implies that genes located at strong TAD boundaries are crucial for coordinating the transcriptional level adjustments of gene expression. In particular, the molecular roles of RNA polymerase II binding and CTD domain binding demonstrate how these genes are involved in both the initiation and completion phases of transcription, which makes them significant for the appropriate control of gene transcription. Important participants in mRNA processing, such as SCAF8 and HNRNPU [73, 74], assure that only correctly processed mRNAs are translated into proteins, hence maintaining cellular effectiveness and functionality.

**Table 14:**
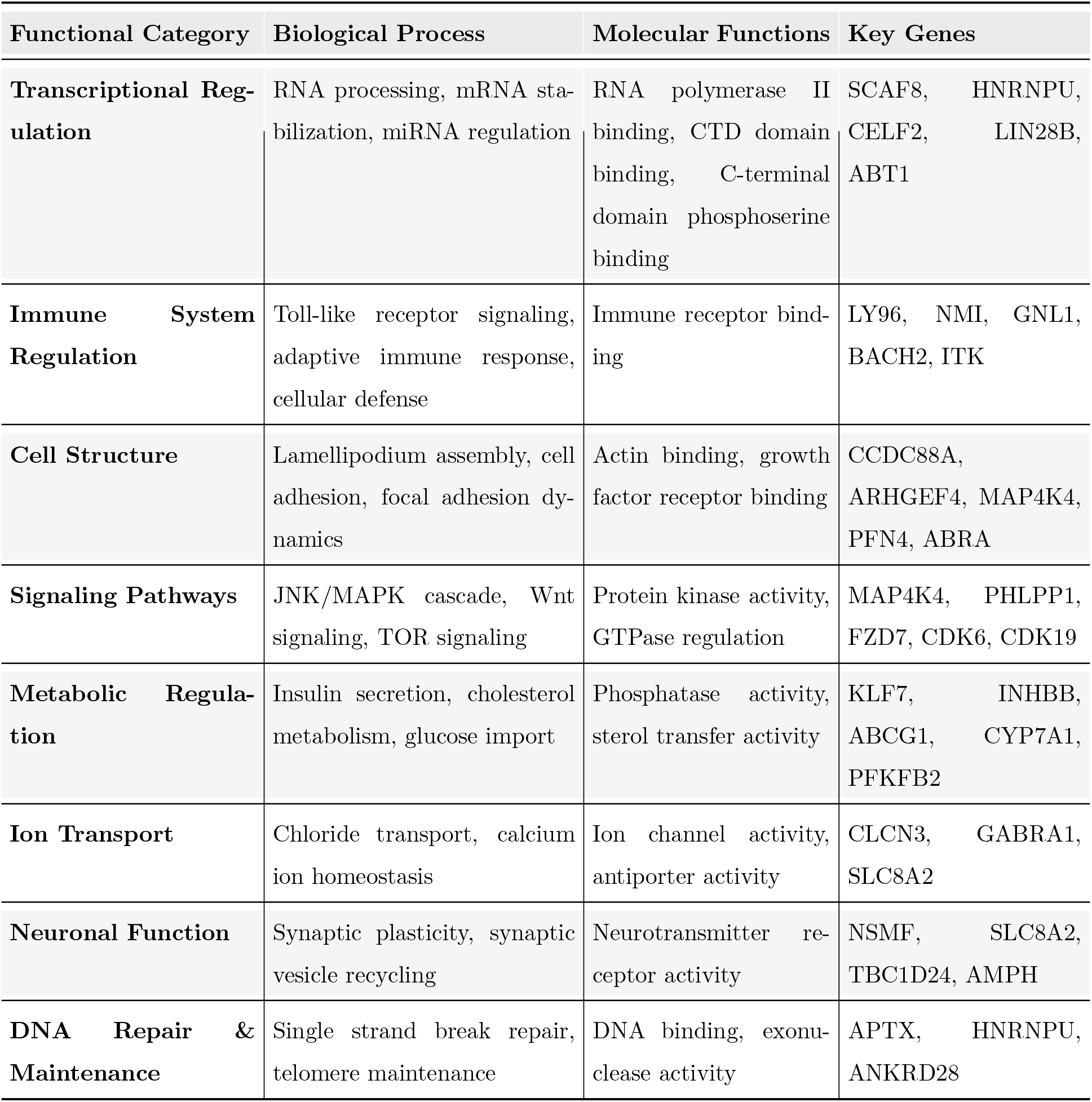
Genes Associated with Strong TAD Boundaries and Their Functional Enrichment.

| Functional Category | Biological Process | Molecular Functions | Key Genes |
| --- | --- | --- | --- |
| **Transcriptional Regulation** | RNA processing, mRNA stabilization, miRNA regulation | RNA polymerase II binding, CTD domain binding, C-terminal domain phosphoserine binding | SCAF8, HNRNPU, CELF2, LIN28B, ABT1 |
| **Immune System Regulation** | Toll-like receptor signaling, adaptive immune response, cellular defense | Immune receptor binding | LY96, NMI, GNL1, BACH2, ITK |
| **Cell Structure** | Lamellipodium assembly, cell adhesion, focal adhesion dynamics | Actin binding, growth factor receptor binding | CCDC88A, ARHGEF4, MAP4K4, PFN4, ABRA |
| **Signaling Pathways** | JNK/MAPK cascade, Wnt signaling, TOR signaling | Protein kinase activity, GTPase regulation | MAP4K4, PHLPP1, FZD7, CDK6, CDK19 |
| **Metabolic Regulation** | Insulin secretion, cholesterol metabolism, glucose import | Phosphatase activity, sterol transfer activity | KLF7, INHBB, ABCG1, CYP7A1, PFKFB2 |
| **Ion Transport** | Chloride transport, calcium ion homeostasis | Ion channel activity, antiporter activity | CLCN3, GABRA1, SLC8A2 |
| **Neuronal Function** | Synaptic plasticity, synaptic vesicle recycling | Neurotransmitter receptor activity | NSMF, SLC8A2, TBC1D24, AMPH |
| **DNA Repair & Maintenance** | Single strand break repair, telomere maintenance | DNA binding, exonuclease activity | APTX, HNRNPU, ANKRD28 |

**Table 15:**
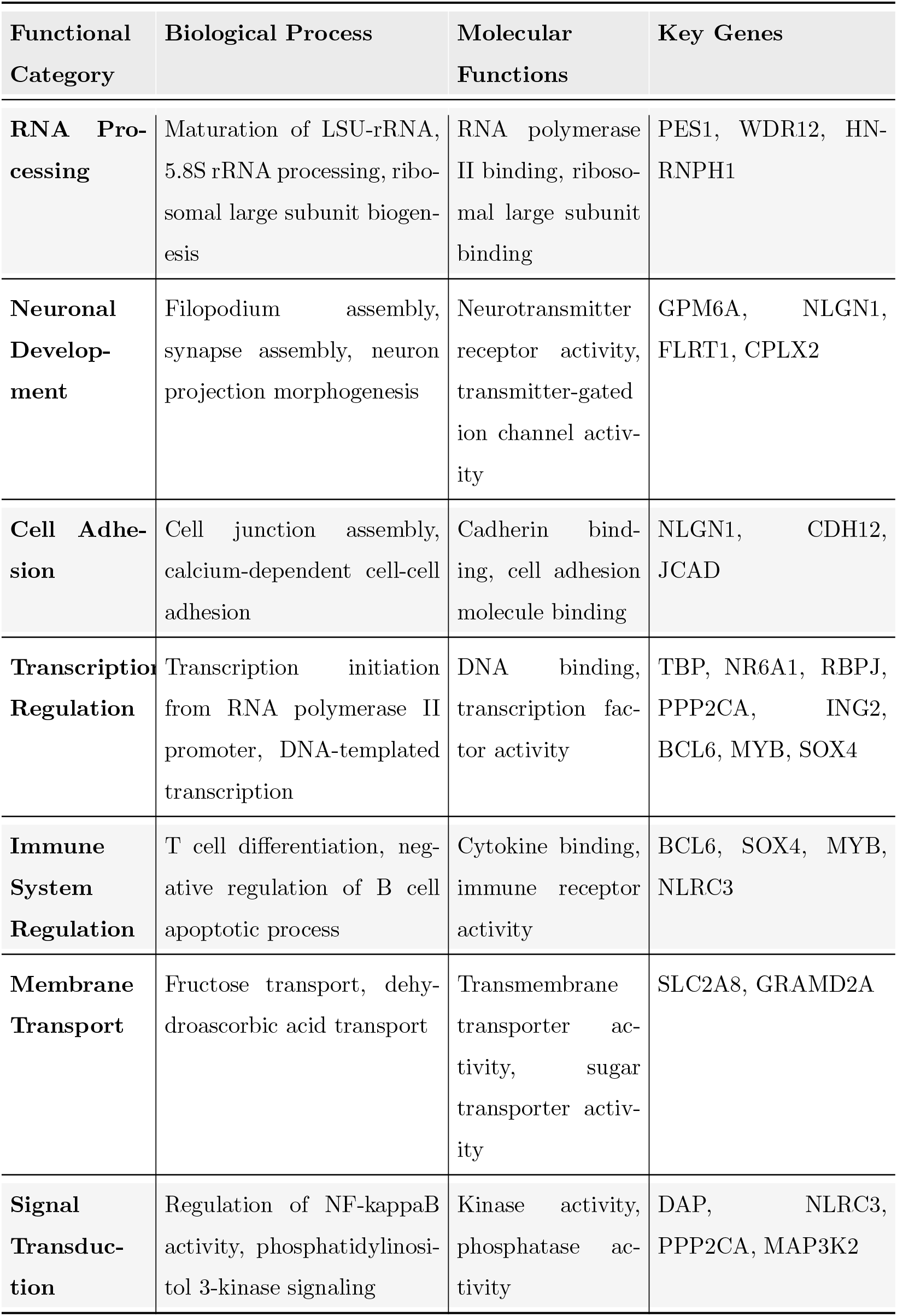
Genes Associated with Weak TAD Boundaries and Their Functional Enrichment.

Lastly, to determine how large-scale chromatin organization might affect the biological activities of genes, we examined the functional enrichment of genes classified by their A/B chromatin compartments. The A compartment’s genes, which usually correspond to open and transcriptionally active regions, are closely linked to ion transport, neuronal development, transcriptional regulation, and signal transduction, suggesting that they serve active roles in basic cellular communication and regulatory mechanisms [75]. In contrast, genes found in the B compartment, which frequently correlate to less transcriptionally active and more compact chromatin, are enriched in processes linked to immune response, metabolic regulation, cell differentiation, and transcriptional control, indicating their role in preserving long-term regulatory programs and specialized cellular states. Detailed categorizations of these functional enrichments can be found in the Supplementary Tables. As a representative example, we highlight the Neuronal Development and Synaptic Function category [76] enriched within the A compartment. This group of genes is enriched for essential biological functions such as synaptic vesicle cycling, neuron projection morphogenesis, filopodium formation, and synapse assembly. These mechanisms underlie neural transmission and connection, both of which are essential for brain development. These genes have a role in synaptic construction and signal transmission at the molecular level through their involvement in PDZ domain binding [77], neurexin binding [78], and syntaxin-1 binding [79]. Their inclusion in the transcriptionally active A compartment emphasizes how genes are spatially regulated to facilitate cellular processes like synapse formation, which are extremely responsive and dynamic.

In conclusion, by examining downregulated gene clusters, the functional properties of genes situated inside strong and weak TAD borders, and the miRNA target genes with the highest and lowest performances, we methodically analyzed the biological implications of our model. The model’s ability to represent biologically significant patterns, such as temporal stability, regulatory architecture, and pathway-specific behaviors, is shown by these examinations. Applying the same functional categorization and enrichment techniques to other miRNA categories, including stable and upregulated genes, allowed us to present a broader perspective (see Supplementary Material).

#### 3.5.2. Ontology-Based Interpretation of mRNA Target Gene Groups

We used the same multi-level categorization approach that we used for miRNA targets to describe the biological relevance of the model’s predictions for the mRNA target gene group. This analysis compares the biological functionality of genes with the highest and lowest performance, groups genes into upregulated, downregulated, and stable categories based on their expression behavior, and examines the biological functions of genes located at strong and weak TAD boundaries and the biological grouping of gene groups located in compartments A and B.

According to an analysis of the highest-performing mRNA target genes, genes such as the core metabolic enzyme HPRT1, which exhibits broad and stable expression [80], as well as VIM and ITGA8, which stand for structural and regulatory factors and typically exhibit an upregulated/stable expression behavior, preserve cytoskeletal integrity and extracellular matrix organization [81]. On the other hand, genes with the lowest performance, such as MCPT4, MMP3, and THTPA, have dynamic values that are specific to particular cell types and even change in response to environmental conditions such as stress and inflammation [82]. At the same time, low-performing genes are typically found in mixed compartmental environments as well as weak boundary regions where regulatory signals are more variable and contextdependent, whereas high-performing genes are typically found in regulatory contexts that support stable and consistent transcription.

Similarly to miRNA, the temporal grouping of mRNA target genes has been divided into Upregulated, Downregulated, and Stable categories based on the slopes of their expression values. Active developmental, immunological, and repair-related functions are indicated by upregulated genes that are enriched in processes such as cell differentiation, extracellular matrix remodeling, and signal transmission. Consistent regulation in vital structural and metabolic pathways is required for stable expression, many of which are crucial for preserving tissue integrity, cell-cell adhesion, and lung-specific homeostasis. Lastly, genes linked to tissue remodeling, fibrosis, and inflammation were downregulated, indicating a decrease in fibrotic or immunological activity.

To investigate the effect of chromatin architecture on gene behavior, we examined the biological behavior of genes located at strong and weak TAD boundaries and genes located between A/B chromatin compartments. Strong boundary genes, such as NMEE3 and Integrin alpha 8 [83], are generally located in stable chromatin regions associated with cell adhesion, growth factor signaling, and structural maintenance in long-lived tissues. On the other hand, weak boundary genes such as IGFBP3 and TGFB1 tend to be located in more dynamic chromatin regions, have variable tissue expression, and are involved in inflammation, tissue remodeling, and immune responses. Compartment B genes are involved in processes like lipid transport, differentiation, and extracellular matrix remodeling, while Compartment A genes are typically involved in metabolic regulation, immune signaling, and cell adhesion, according to our examination of the compartment analysis.

Groupings such as correlation-based sorting, temporal slope-based clustering, and chromatin architecture were used to interpret mRNA target genes biologically and highlight the relationships between biological functionality and model performance. The highest and lowest performing genes show different patterns in areas such as chromatin organization and TAD boundaries, demonstrating the importance of both gene functionality and genome architecture behaviors on predictions. Detailed tables containing the full functional categorizations, enrichment results, and gene specific characteristics for mRNA target genes are provided in the Supplementary Material.

## 4. Conclusion

In this study, we introduced STEPmr and STEPmi, two novel spatiotemporal graph neural network frameworks for predicting temporal mRNA and miRNA expression by integrating high-resolution Hi-C genomic interaction data with additional biologically meaningful features. Our approach addresses the limitations of prior static or purely temporal models by explicitly modeling both the structural organization of the genome and the temporal dynamics of gene regulation. By incorporating temporal Node2vec embeddings, biologically-informed edge weights, and a temporal-aware composite loss function, we were able to capture complex, long-range dependencies in gene expression patterns that emerge from the interplay between 3D genome architecture and regulatory interactions.

Experimental results on publicly available miRNA and mRNA time-series datasets demonstrated that both models achieved improvements in predictive accuracy over baseline spatio-temporal, sequence-based, and recurrent approaches, with STEPmi attaining a mean Pearson correlation of 0.93 and STEPmr reaching 0.77. Importantly, our analysis revealed that the models are particularly effective at predicting genes with stable or broadly conserved expression profiles across tissues, while genes exhibiting highly context-dependent or condition-specific regulation remain more challenging.

Beyond predictive performance, Gene Ontology enrichment analyses provided biological validation for the model outputs. The highest-performing genes were enriched for roles in fundamental processes such as transcriptional regulation, neuronal development, and immune system modulation, often residing within well-defined chromatin compartments. Conversely, lower performing genes were linked to dynamic processes such as tissue remodeling and inflammatory response, reflecting the inherent variability and environmental sensitivity of these expression patterns.

Overall, our work establishes the feasibility and utility of spatio-temporal GNNs in modeling dynamic gene expression and demonstrates that integrating multi-omics structural data with advanced temporal graph learning methods can substantially enhance predictive power and interpretability. The proposed frameworks open new avenues for studying temporal regulation in diverse biological contexts, from developmental processes to disease progression, and could be extended to incorporate other omics modalities, higher-resolution chromatin interaction maps, and cross-species comparative analyses. All code and datasets have been made publicly available, supporting reproducibility and enabling future research to build upon our contributions.

## Declarations

### Availability of data and materials

The datasets used and analyzed during the current study are available on https://github.com/seferlab/temporalgene.

### Code Availability

The proposed method and analysis code is available on https://github.com/seferlab/temporalgene.

### Authors’ contributions

BK: Methodology, Software, Writing. ES: Conceptualization, Methodology, Writing. All authors read and approved the final manuscript.

